# Notch induces transcription by stimulating release of paused RNA Polymerase II

**DOI:** 10.1101/2024.06.13.598853

**Authors:** Julia M Rogers, Claudia A Mimoso, Benjamin JE Martin, Alexandre P Martin, Jon C Aster, Karen Adelman, Stephen C Blacklow

## Abstract

Notch proteins undergo ligand-induced proteolysis to release a nuclear effector that influences a wide range of cellular processes by regulating transcription. Despite years of study, however, how Notch induces the transcription of its target genes remains unclear. Here, we comprehensively examined the response to human Notch1 across a time course of activation using high-resolution genomic assays of chromatin accessibility and nascent RNA production. Our data reveal that Notch induces target gene transcription primarily by releasing paused RNA polymerase II (RNAPII). Moreover, in contrast to prevailing models suggesting that Notch acts by promoting chromatin accessibility, we found that open chromatin was established at Notch-responsive regulatory elements prior to Notch signal induction, through SWI/SNF-mediated remodeling. Together, these studies show that the nuclear response to Notch signaling is dictated by the pre-existing chromatin state and RNAPII distribution at the time of signal activation.

## INTRODUCTION

Cells reliably and precisely convert signaling inputs into cellular decisions by inducing gene expression programs that specify cellular identity. Notch-Delta signaling is one of several essential metazoan pathways of cell-cell communication that guides cell fate decisions in organismal development and homeostasis across tissues and organ systems^1^. Notch signaling is dysregulated in multiple cancer types, including T-cell acute lymphoblastic leukemia, which is associated with gain of function mutations in *NOTCH1*, and squamous cell carcinoma, which is associated with loss of function of *NOTCH1, NOTCH2,* and/or *NOTCH3*^2^.

Notch proteins, the receiver components of this signaling system, are single-pass transmembrane receptors that have an extracellular ligand-binding domain, a juxtamembrane regulatory domain, and an intracellular effector domain (NICD) that is liberated by ligand-induced proteolysis to function as a transcriptional effector^3^. In the nucleus, NICD forms a multiprotein Notch transcriptional complex (NTC) with a DNA-binding transcription factor called RBPJ in mammals (*suppressor of hairless* in flies), and a protein of the Mastermind-like family (MAML). Formation of the NTC leads to transcription of target genes.

Notch signals activate a vastly different array of transcriptional targets in each cell type, allowing this key regulator to have distinct cellular effects depending on the cellular context^3^. As a result, Notch signaling can be oncogenic or tumor suppressive depending on the cancer type^2^. However, how cell type-specificity of Notch targets is achieved and how the NTC stimulates transcription remains unclear at the molecular level. Understanding the molecular mechanisms by which Notch signaling induces transcription is key to understanding how this essential signaling pathway functions in development and disease.

Transcription factors (TFs) can stimulate RNAPII-dependent transcription at multiple steps in the transcription cycle, including during the establishment of chromatin accessibility, RNAPII recruitment to promoters and transcription initiation, or the release of paused RNAPII into productive elongation^4,5^. Previous work has led to conflicting models of how the NTC stimulates transcription. The prevailing model suggests that the NTC cooperates with chromatin remodelers to increase chromatin accessibility^6^. Consistent with this idea, genetic associations between components of the SWI/SNF chromatin remodeler complex and Notch signaling have been observed in both mice and flies^6,7^. Immunoprecipitation followed by mass spectrometry, co-immunoprecipitation, and proximity labeling studies have also suggested that components of the SWI/SNF complex may associate with or are in the molecular neighborhood of Notch1^7–10^. Notably, cells maintained in a persistent Notch-on state have higher chromatin accessibility at Notch binding sites than cells in a Notch-off state^6,11^. Knockdown of SWI/SNF components diminishes this elevated accessibility and reduces transcription of Notch-responsive genes^6,8^.

Other observations are less consistent with a model wherein Notch directly interacts with or recruits SWI/SNF to promote chromatin accessibility. For example, BRM, the catalytic subunit of the SWI/SNF complex, was shown to be present at some Notch responsive promoters prior to Notch activation in mouse cells^10^. Indeed, chromatin compaction can restrict Notch activity, and NTC binding is limited to regions of already accessible chromatin with epigenetic signatures associated with active regulatory regions^12,13^. These findings suggest the possibility of an alternative model, in which gene activation upon Notch signaling relies on cell type-specific transcription factors that establish the proper chromatin context for Notch activation in different cellular contexts^12,14,15^. The fact that Notch signaling induces different programs of gene expression in different cell types^16^ would also be consistent with a model in which the NTC acts on preexisting, poised regulatory regions.

Whether and how interplay between NTC and chromatin remodeling complexes drives specific gene expression upon Notch activation remains poorly understood, largely because studies investigating the dynamics of NTC recruitment and gene activation lack the temporal resolution needed to distinguish direct from indirect effects. We thus sought to elucidate, with high spatial and temporal resolution, how Notch activation induces transcription. We monitored the genomic response to human Notch1 as a function of time after activation in Notch-naïve squamous cell carcinoma (SCC) cells. Newly synthesized RNAs were measured using Transient Transcriptome Sequencing (TT-seq), accessibility was monitored by the Assay for Transposase-Accessible Chromatin with sequencing (ATAC-seq), and active RNAPII was mapped using Precision Run-On Sequencing (PRO-seq). Strikingly, we found that the chromatin accessibility of NTC binding sites did not increase in response to Notch activation. Instead, we found that SWI/SNF establishes accessibility at Notch-responsive regulatory elements prior to signaling, allowing Notch to rapidly bind these loci upon activation. Importantly, we defined the step in the transcription cycle regulated by Notch, finding that the NTC predominantly acts by releasing paused RNAPII into productive elongation at target genes. Together, our data elucidate how cellular context and the preexisting chromatin landscape dictate the specificity of the Notch transcriptional response, providing a conceptual and experimental framework for a better understanding of signal responsive gene expression.

## RESULTS

### Cellular system for time-resolved studies

To study how Notch induces transcription, we used the SC2 squamous cell carcinoma cell line^17^, which is engineered to express a ligand independent, autonomously active form of human Notch1 (ΔEGF-L1596H) that can be silenced by a gamma secretase inhibitor (GSI). These SC2 cells can be maintained in a Notch-naïve state by culturing cells in the presence of GSI. GSI washout then allows us to rapidly toggle cells from a Notch-off to a Notch-on state, thereby enabling the measurement of the direct response of these cells to Notch activation^17^. These features provide distinct advantages for kinetic assessment of how cell state and chromatin context affect the genomic response to a Notch signal.

### Identification of Notch target genes

We identified direct Notch target genes in SC2 cells by performing TT-seq at 1 h and 4 h timepoints after inducing Notch activity by GSI washout (Figure 1A). TT-seq is a metabolic labeling approach used to identify and quantify newly synthesized RNAs^18^, and therefore enables us to define genes induced at specific times after Notch activation. These early time points after Notch activation were chosen to enrich for direct transcriptional responses to Notch activity. To identify specific Notch targets, we included matched time point mock washout control samples, which were subjected to the same manipulations and media change as the GSI washout samples, but GSI was maintained in the media. Notably, most of the genes induced in response to GSI washout were also induced in the mock washout (Figures 1B, S1A), indicating a strong general impact of these cellular manipulations and highlighting the importance of this experimental control.

**Figure 1.**
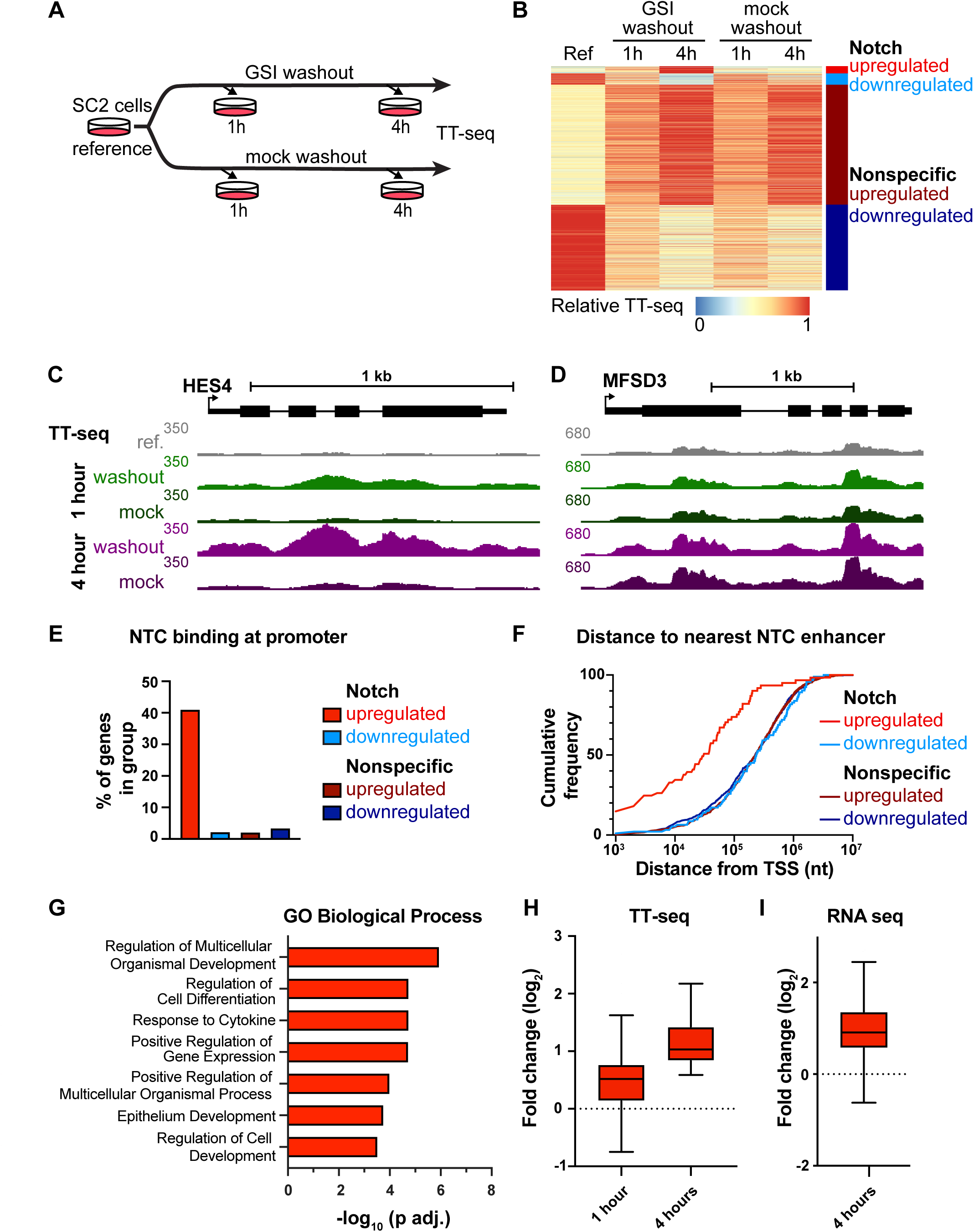
Identification of high-confidence Notch target genes in SCC cells. (A) Design of TT-seq experiment. The reference sample was subjected to mock washout, immediately incubated with labeling media for 10 minutes, and then harvested. The 1 h and 4 h samples were subjected to mock or GSI washout and incubated with labeling media for the 10 minutes preceding harvest at the indicated times. (B) Heatmap indicating relative TT-seq signal over exons for significantly changed genes. Color bars on the right indicate gene groups: Notch upregulated (red, n=61), Notch downregulated (light blue, n=93), nonspecific upregulated (dark red, n=997), and nonspecific downregulated (dark blue, n=711). (C, D) Genome browser images showing sense strand TT-seq reads for a representative Notch upregulated gene (HES4, C) and a representative Nonspecific upregulated gene (MFSD3, D). (E) Percentage of genes with Notch transcription complex (NTC) binding^17^ at the promoter plotted for each gene group. (F) Cumulative distribution plot of the distance from the TSS of genes in each group to the nearest NTC-bound enhancer. (G) Top Gene Ontology – Biological Process enriched terms for genes in the Notch upregulated group. P-values are from the hypergeometric test, corrected for multiple hypothesis according to the Benjamini-Hochberg method. (H) Boxplots showing the fold change (log_2_) in TT-seq exon counts for Notch upregulated genes at 1 and 4 hours, compared to the reference condition. The middle line indicates the median, the box represents the 25^th^ and 75^th^ percentile, and whiskers show 1.5x the interquartile range, or the largest or smallest value in the data set (Tukey method). (I) Boxplots (rendered as in H) showing the fold change (log_2_) in RNA-seq gene counts for Notch upregulated genes relative to the mock washout condition at 4 h. RNA-seq data is from Pan et al.^17^ See also Supplemental Figure S1.

We classified gene responses into four categories to separate Notch-dependent transcriptional responses from Notch-independent ones: Notch-upregulated (upregulated in GSI washout but not mock washout, n=61), Notch-downregulated (downregulated in GSI washout but not mock washout, n=93), nonspecific-upregulated (upregulated in both GSI and mock washout, n= 997), and nonspecific-downregulated (downregulated in both GSI and mock washout, n= 711) (Table S1). As anticipated, the Notch-upregulated gene HES4 was specifically induced under GSI washout but not mock washout conditions (Figure 1C), whereas a representative nonspecific-upregulated gene (MFSD3) was activated in both conditions (Figure 1D).

To further validate that the Notch-upregulated genes are direct targets of Notch activity, we analyzed genomic binding by Notch transcription complex (NTC) components in SC2 cells^17^. For this analysis, we defined NTC binding sites as those loci exhibiting both RBPJ and MAML1 binding by ChIP-seq 4 h after GSI washout. We found that Notch-upregulated genes are much more likely to have NTC binding sites within 1 kb of their transcription start sites, and that these genes are much closer to NTC-bound enhancers than genes within the three other groups (Figures 1E,F), consistent with direct regulation of these genes in response to NTC binding. In contrast, the lack of NTC binding near genes that are downregulated upon GSI washout suggests that gene repression in response to Notch activation is not mediated directly by NTC binding. This repression may instead result from secondary effects, such as Notch induction of transcriptional repressors like the HES genes. The Notch-upregulated gene set contained canonical Notch targets upregulated in many cell types (HES1, HES4, NRARP), known Notch targets in squamous cells (IER5, RHOV), and targets not previously associated with Notch signaling (CELSR2, ADGRG6). The biological process GO terms for this group of genes included terms associated with development and differentiation, consistent with the role of Notch signaling in determining cell fate (Figure 1G). In contrast, the genes whose expression changed in response to mock washout are enriched for GO terms associated with metabolism and biosynthesis, consistent with a response to the replenishment of nutrients in the fresh media used during washout (Figures S1C,D).

While some Notch-upregulated genes are induced within 1 h of GSI washout, they showed maximal expression at 4 h after GSI washout (Figure 1H). Consistent with the increase in RNA synthesis observed in TT-seq, these genes were also induced in published steady-state RNA-seq performed in these cells following 4 h of GSI washout^17^ (Figure 1I).

### Notch activates gene expression without increasing chromatin accessibility

Prior studies have found that Notch binding sites in “Notch-on” cells have higher chromatin accessibility at steady state than in “Notch-off” cells^6,11^. We therefore tested whether Notch directly affects the chromatin landscape by monitoring dynamic changes in chromatin accessibility at time points up to 4 h after Notch induction by GSI washout using ATAC-seq (Figures 2A, S2)^19,20^. Strikingly, Notch-upregulated genes have accessible promoters prior to Notch induction, based on the significant levels of promoter-proximal ATAC-seq reads observed in the reference condition (Figure 2B). This observation highlights how the basal context of a cell, likely defined by other transcription factors, can determine where the NTC can induce gene activity.

**Figure 2.**
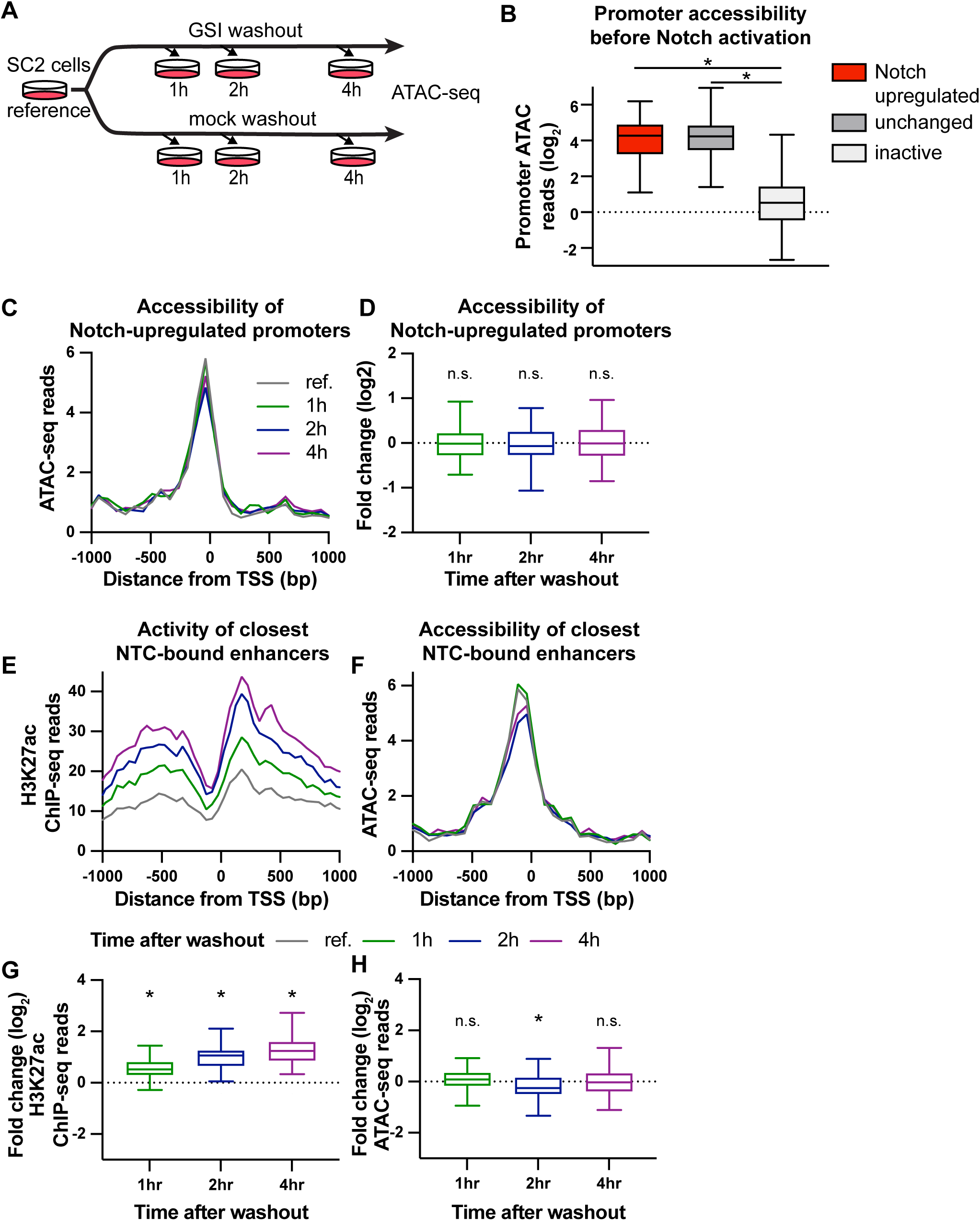
Notch activation does not increase chromatin accessibility. (A) Design of ATAC-seq experiment. The reference condition was subjected to mock washout, and then immediately harvested. All other samples were harvested at the indicated time after GSI washout. (B) Boxplots (rendered as in Figure 1H) showing ATAC-seq reads (log_2_) over the promoters (TSS-450 to TSS+149) of Notch-upregulated genes (n=61), unchanged genes (n=51484) and inactive genes (n=1295). Unchanged genes are defined as genes with a TT-seq gene body read |FC| < 1.1 in the 1h washout and 4h washout condition compared to the reference. Lowly active genes (inactive) are defined as genes with fewer than 10 PRO-seq reads within the promoter region (TSS to TSS+150) at any timepoint. Stars indicate significant differences (p. adj < 0.05) in accessibility by Dunn’s multiple comparisons test. (C) Aggregate plot showing ATAC-seq reads around promoters of Notch-upregulated genes (n=61). ATAC-seq reads are shown in 75 bp bins. (D) Boxplots (rendered as in B) showing fold change (log_2_) in ATAC-seq reads at the promoter (TSS-450 to TSS+149) for Notch-upregulated genes (n=61). “n.s.” indicates conditions not significantly different (p. adj > 0.05) from a fold change of 0 (Wilcoxon signed rank test, p values corrected according to the Benjamini-Hochberg method). (E-F) Aggregate plots showing H3K27ac ChIP-seq (E), or ATAC-seq (F) reads at closest NTC-bound enhancers with identified TSSs (see methods) to Notch-upregulated genes (n=56). E is shown in 50 bp bins, F is in 75 bp bins. (G-H) Boxplots (rendered as in B) showing fold change (log_2_) in H3K27ac ChIP-seq (G), or ATAC-seq (H) reads at closest NTC-bound enhancers to Notch-upregulated genes (n=58). Stars indicate conditions significantly different (p. adj < 0.05) from a fold change of 0 (Wilcoxon signed rank test, p values corrected according to the Benjamini-Hochberg method). See also Supplemental Figure S2.

We next investigated whether Notch activation affects the accessibility of responsive promoters. Aggregate plots around gene promoters and counting of the total ATAC-seq reads over the promoters of Notch-upregulated genes showed no changes in promoter accessibility over the 4 h time course (Figures 2C,D) despite clear increases in gene expression. These observations are inconsistent with a model in which Notch recruits SWI/SNF complexes to open up target gene promoters, instead suggesting that NTCs act on loci that are accessible prior to Notch activation.

Because most Notch genomic binding occurs at distal regulatory regions in other cell types^21^, we also examined whether Notch activation affects the accessibility of Notch-responsive enhancers. We defined NTC-bound regions as those occupied by both NTC subunits RBPJ and MAML1 in published ChIP-seq data ^17^, and used these data to identify the closest NTC-bound enhancer to each Notch-upregulated gene (see methods). We then examined the activity of these enhancers in a 4-hour time course after Notch induction by GSI washout. Enhancer activity, as judged by H3K27ac chromatin immunoprecipitation followed by sequencing (ChIP-seq) (Figures 2E,G), increased over the 4 h time course, confirming that these elements are high-confidence Notch-responsive enhancers. However, these NTC-bound enhancers did not show increased accessibility following GSI washout, either in aggregate plots or in read counts over the enhancers, in agreement with our findings at Notch-responsive promoters (Figures 2F,H). Together, these findings argue against models wherein Notch activation induces gene expression through modulation of chromatin accessibility at either promoters or enhancers. Instead, our data favor a mechanism in which Notch binding occurs at pre-existing open regulatory regions.

### SWI/SNF activity is required to maintain promoter accessibility

Although chromatin accessibility was not increased at sites of Notch-dependent gene induction, previous studies have shown strong associations between the SWI/SNF chromatin remodeling complex and Notch activity^6,7^. To elucidate the basis for the interplay between SWI/SNF and Notch, we performed TT-seq to examine newly synthesized transcripts at 1 and 4 h time points after Notch activation in the presence of an allosteric-inhibitor of SWI/SNF activity, BRM014, which rapidly inactivates the SWI/SNF ATPase, resulting in decreased promoter and enhancer accessibility^22–25^ (Figures S3A,B). Adding BRM014 at the time of Notch activation blocked the induction of 41 of the 61 Notch-upregulated genes (Figure 3A), consistent with previous work linking SWI/SNF activity to Notch-dependent gene activation. The chromatin accessibility in the reference condition was not different between SWI/SNF dependent or independent gene groups (Figure 3B), nor did either subgroup show any change in accessibility after Notch induction (Figure 3C). As expected, SWI/SNF independent genes did not exhibit a significant change in accessibility as assessed by ATAC-seq following BRM014 treatment (Figures 3D,F). However, adding BRM014 at the moment of GSI or mock washout revealed a rapid and significant decrease in ATAC-seq signal at SWI/SNF dependent promoters (Figures 3E,F). Importantly, because loss of accessibility upon BRM014 treatment was observed under mock washout conditions, the sensitivity of promoter accessibility to SWI/SNF appears to be independent of the Notch activation status of the cells. These data suggest that these SWI/SNF dependent promoters have a persistent reliance on SWI/SNF remodeling to remain accessible, regardless of whether Notch is active. Suppression of SWI/SNF activity, even at the moment of GSI washout, is sufficient to reduce chromatin accessibility and prevent Notch-mediated gene activation.

**Figure 3.**
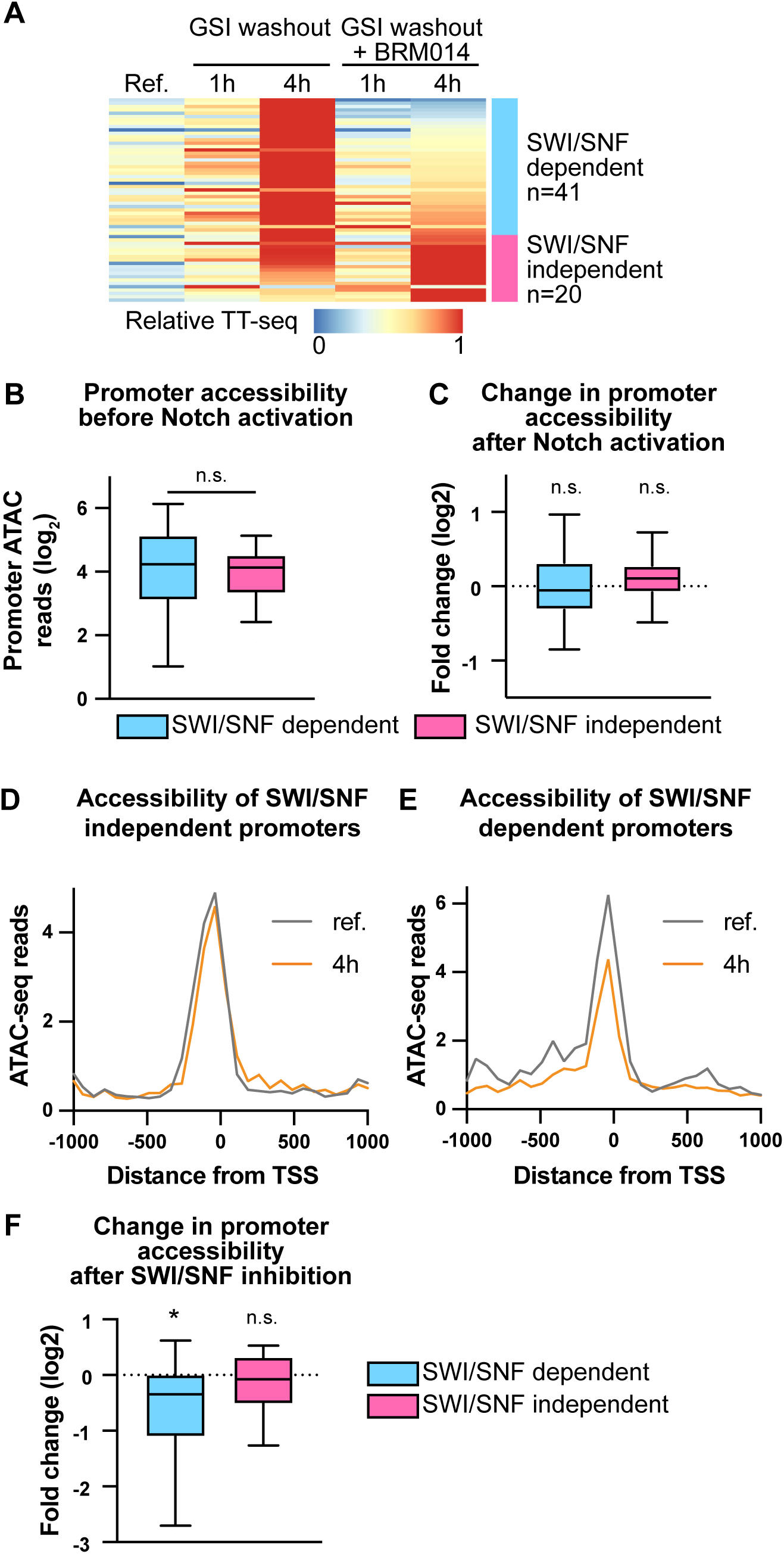
SWI/SNF activity is required to keep most promoters accessible for activated Notch. (A) Heatmap indicating relative TT-seq signal over exons for Notch-upregulated genes. Color bars on the right indicate gene groups: SWI/SNF dependent (blue, n=41), SWI/SNF independent (pink, n=20). (B) Boxplots (rendered as in Figure 1H) showing ATAC-seq reads (log_2_) over the promoters (TSS-450 to TSS+149) of SWI/SNF dependent and SWI/SNF independent genes. “n.s.” indicates groups not significantly different (p > 0.05) by Mann-Whitney test. (C) Boxplots (rendered as in B) showing fold change (log_2_) in ATAC-seq reads at the promoter (TSS-450 to TSS+149) for SWI/SNF dependent and SWI/SNF independent genes at 4 hours after Notch activation compared to the reference condition. “n.s.” indicates conditions not significantly different (p > 0.05) from a fold change of 0 (Wilcoxon signed rank test). (D-E) Aggregate plot showing ATAC-seq reads around promoters of SWI/SNF independent (D) and SWI/SNF dependent (E) genes. ATAC-seq reads are shown in 75 bp bins. (F) Boxplots (rendered as in B) showing fold change (log_2_) in ATAC-seq reads at the promoter (TSS-450 to TSS+149) for SWI/SNF dependent and SWI/SNF independent genes at 4 hours after mock washout in the presence of BRM014, compared to the reference condition. Stars indicates conditions significantly different (p < 0.05) from a fold change of 0 (Wilcoxon signed rank test). See also Figure S3.

### Notch-upregulated genes are activated by release of paused RNAPII

If Notch is not acting to increase chromatin accessibility, how does it stimulate transcription? To address this question, we performed PRO-seq^26,27^ to measure the effect of activated Notch on RNAPII at a series of time points from 15 minutes up to four hours after GSI washout (Figures 4A, S4A). PRO-seq maps the location of engaged RNAPII across the genome, which allows us to measure both gene body transcription as well as RNAPII occupancy at promoters. Notch-upregulated genes showed an increase in PRO-seq signal in the gene body within 30 minutes of Notch activation (Figure 4B) that persists over the time course, consistent with direct upregulation of these genes by increased transcription. If Notch activation served to increase the recruitment of RNAPII or transcription initiation, we should observe an increase in PRO-seq signal at Notch-upregulated promoters over this time window. However, the promoter PRO-seq signal did not increase upon Notch activation and was instead significantly decreased at 30 min and 1 h after GSI washout (Figure 4C). We conclude that Notch activation does not primarily increase transcription by stimulating initiation.

**Figure 4.**
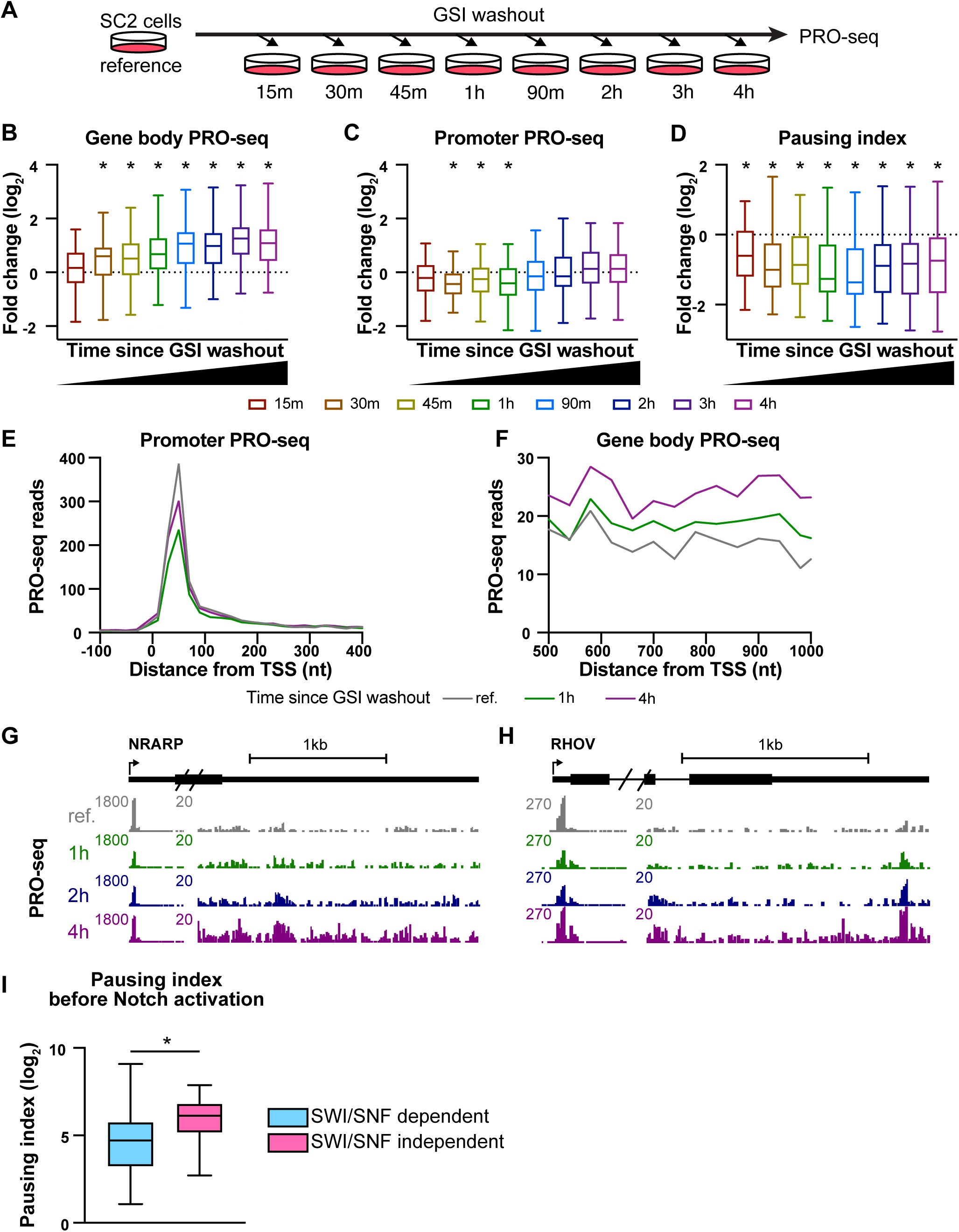
Notch-dependent genes are activated by pause release. (A) Design of PRO-seq experiment. The reference condition was subjected to mock washout, and then immediately harvested. All other samples were harvested at the indicated time after GSI washout. (B,C) Boxplots (rendered as in Figure 1H) showing the fold change (log_2_) in PRO-seq gene body (B) or promoter (C) counts for Notch upregulated genes, compared to the reference condition. Gene body windows are defined as TSS + 250 bp to TSS + 5 kb, and promoter windows are defined as TSS to TSS + 150 bp. Stars indicate conditions significantly different (p. adj < 0.05) from a fold change of 0 (Wilcoxon signed rank test, p values corrected according to the Benjamini-Hochberg method). (D) Boxplots (rendered as in Figure 1H) showing the fold change (log_2_) in pausing index compared to the reference condition for Notch upregulated genes. Pausing index is defined as promoter counts per kb / gene body counts per kb. Stars indicate conditions significantly different (p. adj < 0.05) from fold change of 0 (Wilcoxon signed rank test, p values corrected according to the Benjamini-Hochberg method). (E) Aggregate plots showing average reads of PRO-seq signal around promoters of Notch upregulated genes. Data are plotted in 20 bp bins. (F) Aggregate plots showing average reads of PRO-seq signal in the gene body of Notch upregulated genes. Data are plotted in 40 bp bins. (G,H) Genome browser images showing PRO-seq reads for Notch upregulated genes NRARP (G) and RHOV (H). The scale of the browser images is reset at 500 bp downstream of the TSS to allow visualization of the gene body signal. (I) Boxplots (rendered as in B) showing pausing index of SWI/SNF dependent and SWI/SNF independent genes in the reference condition. Stars indicate groups significantly different (p < 0.05) by Mann-Whitney test. See also Supplemental Figure S4.

Transcription can be induced by stimulating transcription initiation or promoting the release of paused RNAPII. To evaluate if Notch activation impacted pause release, we calculated the pausing index for the Notch-upregulated genes over time. The pausing index, a ratio of promoter to gene body PRO-seq reads, indicates the level of RNAPII pausing for each gene. We found that the pausing index decreased significantly at Notch upregulated genes as early as 15 min after Notch activation (Figure 4D), implicating a pause release mechanism in Notch-dependent gene activation.

Aggregate plots showing the average signal around promoters and gene bodies of the Notch-upregulated genes further support the presence of a pause release mechanism (Figure 4E,F). At 1 h after Notch activation, there were fewer PRO-seq reads at the promoter and more in the gene body, indicative of the transition of RNAPII from pausing to productive elongation. By 4h, the gene body signal increased further, and the promoter signal rebounded to approach that of the reference condition, consistent with re-initiation after the initial release of paused RNAPII.

Browser shots of two Notch regulated genes, NRARP (Figure 4G) and RHOV (Figure 4H) further highlight the effect of Notch activation in stimulating pause release. In the reference condition (*i.e.* before Notch activation), both promoters exhibit a peak of paused RNAPII proximal to the promoter with very low signal in the gene body. At 1 h, a decrease in the paused peak is accompanied by release of RNAPII into the gene body, with highest gene body signal observed at 4 h. There is also restoration of the PRO-seq signal at the promoter at 4 h, consistent with increased transcriptional initiation by RNAPII at this time point. This delayed stimulation of initiation is particularly evident for the canonical Notch responsive gene HES1, which responds rapidly to Notch activation and shows oscillations in PRO-seq signal over time, with a period of roughly 2 h, consistent with previous studies^28^ (Figure S4B).

Because RNAPII pausing has been associated with maintaining promoter accessibility^29^, we asked whether pausing itself could explain the SWI/SNF independence of some of the Notch responsive genes. We found that the SWI/SNF independent genes exhibited a significantly higher pausing index than genes that were sensitive to acute SWI/SNF inhibition (Figure 4I). Therefore, we propose that there are two mechanisms by which promoters can be maintained in a Notch-responsive state (Figure 5). For most responsive genes, SWI/SNF is required both to establish chromatin accessibility and to maintain promoters in a Notch-responsive state, allowing direct binding of NTC to either the promoter or enhancer to promote RNAPII pause release. For a smaller proportion of genes, high levels of stably paused RNAPII can maintain promoter accessibility in the absence of SWI/SNF activity, allowing access by NTC upon Notch pathway activation, even when SWI/SNF is inhibited.

**Figure 5.**
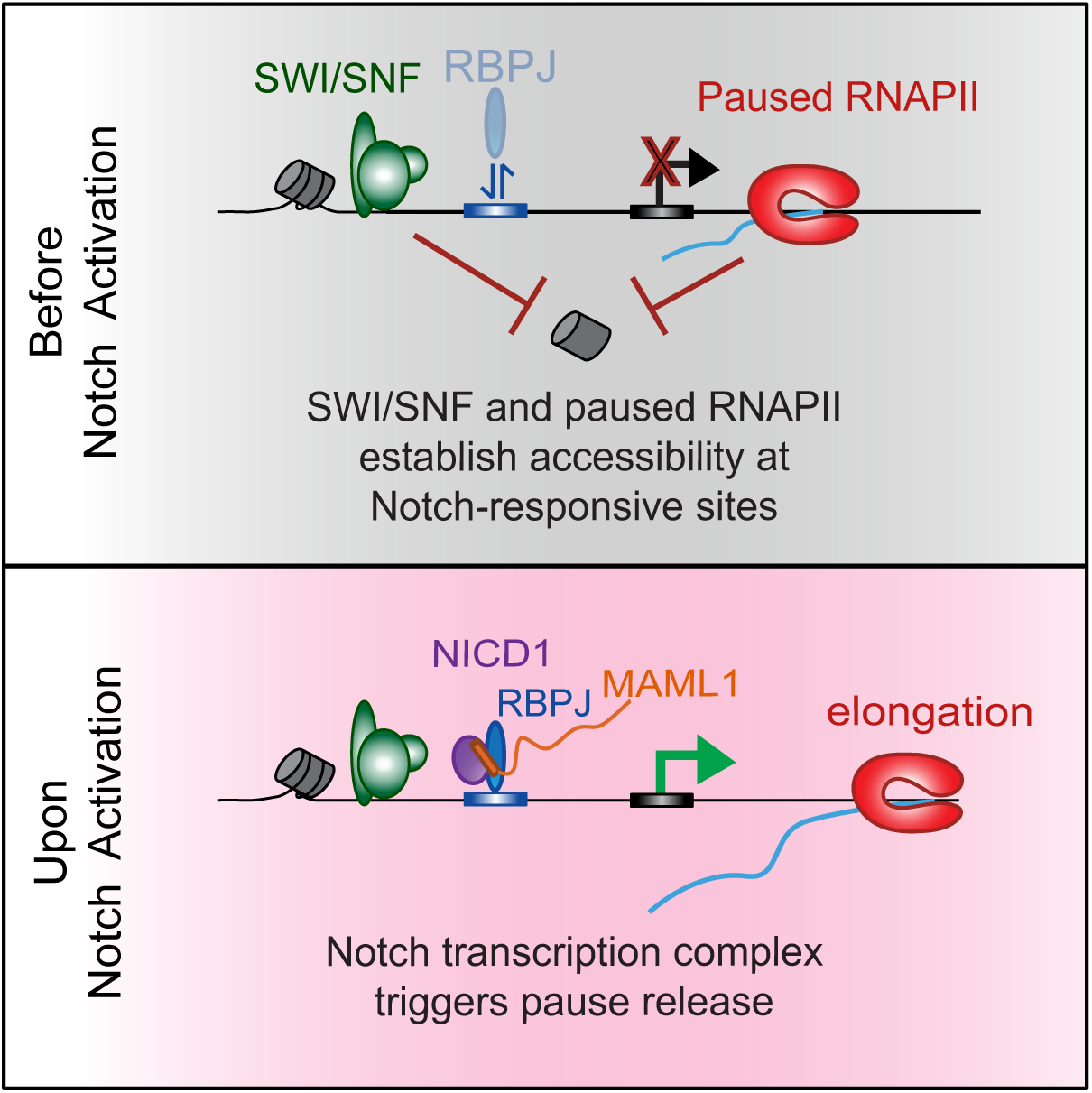
Model figure depicting pause release mechanism for NTC-induced transcription.

## DISCUSSION

Using time-resolved genomic approaches in Notch-naïve SC2 cells, we found that activated Notch stimulates gene activity primarily by promoting release of paused RNAPII. Our data also revealed that NTCs activate pre-existing accessible sites rather than by inducing regions of chromatin accessibility. These findings clarify the associations between SWI/SNF and Notch activation, demonstrating the importance of SWI/SNF for establishing, and in most cases maintaining, the permissive chromatin state needed for Notch-dependent gene induction. The reliance of NTCs on the prior opening of chromatin by other transcription factors rationalizes why Notch signaling is able to induce expression of different target genes in different cell types and underscores the cooperation of Notch nuclear complexes with distinct TFs in each cell type. Our findings dovetail nicely with other systems in which signal-responsive and developmentally regulated expression programs are coordinated by combinations of TFs, with lineage-determining TFs establishing the genomic landscape of accessible regulatory elements that are available for binding by signal-responsive TFs^30–32^. We speculate that other signal-dependent transcription networks function similarly, with signal-responsive TFs themselves playing little role in defining chromatin structure. Instead, chromatin remodelers collaborate with other TFs, prior to signal induction, to generate the appropriate chromatin landscape for cell-type or condition specific binding of signal-responsive TFs. Indeed, like NTCs, NF-kB also binds to previously accessible chromatin^33–35^ and has been implicated in regulating transcription elongation and pause release^36,37^, highlighting parallels between Notch signaling and inflammatory pathways.

Several features of our experimental design – synchronization of Notch activation by GSI washout, time-resolved analyses of gene induction, readouts of nascent transcription, and rapid acting SWI/SNF inhibitors – made it possible to observe direct responses to Notch signal activation and draw mechanistic inferences from these studies. Notably, the rigorous setup of our system allowed us to observe a substantial effect of media change, apparent in our mock-washout matched time point controls. This finding raises a concern that should be taken into account by the gene expression field anytime that media washout or exchange is used to identify transcriptional targets (e.g., when GSI washout is used to identify Notch-induced genes). Accordingly, by rigorously filtering out non-specific effects, we identified fewer Notch target genes than described in some studies, but find the genes identified to be high confidence, direct targets. Indeed, the number of Notch-regulated genes studied here is comparable to that seen in studies identifying Notch targets in T-cell acute lymphoblastic leukemia/lymphoma^38^.

Why might a pause release mechanism be advantageous for the action of Notch nuclear complexes? RNAPII pausing is known to promotes synchronicity in gene activation, for example in the context of developing fly embryos to ensure coordinated cell behavior in tissue development^39,40^. Because Notch signals act at discrete times during tissue development to direct cell fate decisions, the stimulation of RNAPII pause release by Notch can direct precise timing of responses and specify robustness in cell fate choices in response to a Notch signal. One particular developmental process that might benefit from the pause release mechanism is somitogenesis, where Notch signals in the segmentation clock are highly synchronous. Interestingly, canonical Notch target genes, including HES1, HES2, and HES4, fall into the SWI/SNF-independent gene group with higher levels of paused RNAPII, consistent with these core targets being primed for robust and synchronous Notch responsiveness.

In conclusion, our data supports a new pause release model for Notch-mediated gene activation. This model establishes the groundwork for future inquiry into the identity of transcription factors responsible for establishing the landscape of paused RNAPII that Notch can act on in distinct cell types, and the basis of how NTC binding promotes RNAPII release. Because the interaction of the MAML1 subunit of NTCs with p300 is required for Notch-dependent gene activation^41,42^, it is possible that pause release is stimulated by histone acetylation, coupled to binding of BRD4 to acetylated histone tails and the stimulation of P-TEFb activity at Notch-responsive promoters^5^. Additionally, it has been reported that the NTC can interact with components of the super elongation complex (SEC) in flies, and could potentially directly recruit P-TEFb to its target genes^43^. It will be interesting in future work to identify the mechanism by which Notch promotes pause release, and to determine whether these partners of Notch are conserved or distinct across various tissues and Notch-associated cancers.

## METHODS

### Data and code availability

TT-seq, PRO-seq, H3K27ac ChIP-seq, and ATAC-seq data sets have been deposited in GEO (GSE269128) and are publicly available upon publication. Custom scripts have been deposited in Zenodo and are publicly available at https://zenodo.org/records/11222086 and https://zenodo.org/records/6654472. Any additional information required to reanalyze the data reported in this paper is available from the lead contact upon request.

### Cell culture and washout protocol

SC2 squamous cell carcinoma cells^17^ were cultured in keratinocyte media (see Supplemental Methods). GSI was prepared as a 1 mM stock of Compound E (EMD Millipore 565790) in DMSO (Sigma-Aldrich 472301). Cells were split every 3-4 days, using 0.25% trypsin (VWR 45000-664) supplemented with 1 μM GSI. Cells were incubated in 0.25% trypsin at 37 °C until cells began to slough off the plate, then were quenched with fresh media and replated. Cells were routinely tested for mycoplasma, and used for experiments within 1 month of thawing a fresh vial.

For all gamma secretase inhibitor (GSI) washouts, media was removed, cells were washed twice with media containing 0.1% DMSO instead of GSI, and replenished with media containing 0.1% DMSO instead of GSI, for a total of three media changes. In the mock washouts, both the washing media and the replenishing media contained 1 μM GSI. For experiments with BRM014 (MedChem Express HY-119374), the replenishing media, but not the washout media, either contained 1µM BRM014 or 0.01% DMSO depending on the sample.

### TT-seq sample preparation and library construction

The evening before the time course, 11 15-cm dishes were seeded with 10 million SC2 cells per plate in SC2 media with 1 μM GSI. 10 plates were used for sample conditions, the 11^th^ plate was used as a sentinel plate: cells were harvested from this plate using 0.25% trypsin and were counted to determine the total number of cells per plate. For each replicate, it was assumed that all 10 experimental plates had the same number of cells as the sentinel plate. Replicates were performed on different days (n=2).

Washout and mock washouts were performed as described above in the GSI washout section. 1 µM BRM014 or the equivalent volume of DMSO was added in the replenishing media after the two media changes. Labeling was performed for 10 minutes in 15 mL of media supplemented with 500 µM 4-Thiouridine (4-SU) (Thermo Fisher J60679) with GSI and/or BRM014 depending on the experimental condition. For the 0 h timepoints, labeling media was added immediately after the mock washouts. For the 1 hr and 4 hr timepoints, labeling media was added at 50 min or 3 h and 50 min after washout, respectively. To harvest cells for TT-seq, plates were quickly rinsed with 20 mL PBS; then 2 mL of Trizol (Thermo Scientific 15596026) was added to the plate. After 3 min of lysis in the dish, the cell lysate in Trizol was collected and frozen at - 80°C. 1.4 mL of each sample was used to prepare RNA. For normalization purposes, fly spike-in cells were used. RNA was then purified and used as input for TT-seq library construction (see Supplemental Methods). Libraries were pooled and sequenced paired-end with the 200 cycle kit on a Novaseq S4 single lane.

### ChIP-seq sample preparation and library construction

Two days before the time course, cells were plated 1:4 into three 15 cm dishes per sample in SC2 media with 1 μM GSI. Replicates were performed on different days (n=2). Cells were crosslinked with 1% formaldehyde at the indicated time post GSI washout, and chromatin was prepared from the cells (see Supplemental Methods). 30 μL H3K27ac antibody (Active Motive 39133) was used for each IP (see Supplementary Methods). For spike normalization, the same amount of sheared Drosophila DNA was added to each sample before library construction. Libraries were prepared using the NEBNext Ultra II DNA Library Prep Kit for Illumina (New England Biolabs E7645) and sequenced paired end on a NovaSeq with an S1 single lane 100 cycle kit.

### ATAC-seq sample preparation and library construction

The evening before the time course, cells were plated in 12 well dishes, at 125,000 cells / well in SC2 media with 1 μM GSI. Replicates were performed on different days (n=2). At the time point after washout or mock washout, cells were rinsed in PBS, and harvested using 0.25 % trypsin with GSI and/or BRM014 according to the sample, and quenched with SC2 media containing GSI and/or BRM014. Cells were counted, and 100,000 cells were transferred to an Eppendorf tube, and combined with 10,000 fly spike-in cells. The spike-in S2 cells had been collected, resuspended in BAMBANKER cryopreservative media (VWR 101974-112), aliquoted, and stored at −80°C. All spike in aliquots for one experimental time course replicate were thawed and combined before beginning the time course.

ATAC was performed according to the OmniATAC protocol^44^ (see Supplemental Methods). Libraries were sequenced paired end on a NovaSeq with an S1 single lane 100 cycle kit.

### PRO-seq sample preparation

The evening before the time course, cells were plated on 9 10-cm dishes, at 5.5 million cells / plate in SC2 media with 1 μM GSI. Replicates were performed on different days (n=2, n=3 for the reference condition). At the relevant time post washout, plates were rinsed with PBS, and 0.25 % trypsin (containing GSI for the mock washout, or an equivalent amount of DMSO for all other time points) was added. Once cells were detaching from the plate, trypsin was quenched with cold SC2 media (containing GSI for the mock washout, or an equivalent amount of DMSO for all other time points), and cells were transferred to a 15 mL falcon tube. Cells were permeabilized, and PRO-seq libraries were constructed as described in Supplemental Methods. Pooled libraries were sequenced using the Illumina 200 cycle kit on an S4 single lane, followed by additional run on a full SP lane for more depth for some samples.

### TT-seq mapping

Reads were mapped first to the spike genome (dm6) using bowtie 1.2.2^45^, then to the human genome (hg38) using STAR2.7.3a^46^. See Supplemental Methods for additional mapping parameters. To normalize samples, reads mapping to exons in the active gene models (see below) were counted using featurecounts, and used to calculate size factors using DESeq2. PCA plots were generated using DESeq2 over the top 500 genes. These size factors were scaled so the minimum was 1 and were used to normalize bedGraphs with the custom script normalize_bedGraph.pl. Biological replicates (n=2) were merged using the script bedgraphs2stbedGraph.pl.

### ChIP-seq mapping

Reads were mapped first to the spike genome (dm6), then to the human genome (hg38) using bowtie1.2.2^45^ with options -k1 -v2 – best. The custom script bowtie2stdBedGraph.pl was used to make bedGraph files with a 75bp shift. As spike returns were not different between time points, samples were depth normalized using the script normalize_bedGraph.pl. Biological replicates (n=2) were merged using the script bedgraphs2stbedGraph.pl. See Supplemental Methods for additional mapping parameters.

### ATAC-seq mapping

Reads were first mapped to the spike genome (dm6) and reads that did not map to the spike were mapped to the human genome (hg38) using bowtie1.2.2^45^. See Supplemental Methods for additional mapping parameters. A custom script, extract_fragments.pl, was used to filter and retain unique reads between 10 and 150 base pairs, which correspond to regions of open chromatin, and convert files into bedGraph format. Spike return rates were not significantly different between samples. To normalize samples, reads mapping to the −1 kb to +1 kb region around active promoters in this cell type (see Genome Annotation below) were counted using the custom script makeheatmap with the “-b v -v t -s b” options. The number of these reads was used to calculate size factors using DESeq2^47^. These size factors were scaled so the minimum was 1, and used to normalize bedGraphs with the custom script normalize_bedGraph.pl. PCA plots were generated using DESeq2 over the top 500 promoters. Biological replicates (n=2) were merged using the script bedgraphs2stbedGraph.pl.

### PRO-seq mapping

Reads were mapped to a combined genome including both the spike (dm6) and primary (hg38) genomes using bowtie2^48^. See Supplemental Methods for additional mapping parameters. As spike percentages were not significantly different between samples, biological replicates were depth normalized using the script normalize_bedGraph.pl. PCA plots were generated using DESeq2 using gene body PRO-seq counts over the top 500 genes. Biological replicates (n=3 for t=0, n=2 for all other time points) were merged using the script bedgraphs2stbedGraph.pl. The merged t=0 bedgraph was normalized by multiplying by 0.66 to using normalize_bedGraph.pl so that all time points had an equivalent depth. BedGraph files were binned in 10 bp intervals, and converted to bigwig files for visualization on the genome browser using UCSCtools^49^.

### RNA-seq mapping

RNA-seq fastq files from Pan et al.^17^ for samples 4 hours after GSI washout (GSM4732270, GSM4732271, GSM4732272) and after mock washout (GSM4732261, GSM4732262, GSM4732263) were downloaded from the sequence read archive. Reads were mapped to hg38 using STAR version 2.7.3a^46^ as described in the TT-seq mapping section. Gene counts were determined using the featurecounts function in the Rsubread^50^ package version 2.14.2, and log_2_ fold change after 4 hours of GSI washout was determined using DESeq2^47^ version 1.40.2. Samples were normalized using the size factors calculated by DESeq2.

### Genome Annotation

To select gene-level features for differential expression analysis, and for pairing with PRO-seq data, we assigned a single, dominant TSS and transcription end site (TES) to each active gene. This was accomplished using a custom script, get_gene_annotations.sh, which uses RNA-seq read abundance and PRO-seq R2 reads (RNA 5’ ends) to identify dominant TSSs, and RNA-seq profiles to define most commonly used TESs. RNA-seq from Pan et al.^17^ at 0hr (GSM4732261, GSM4732262, GSM4732263) and 4hr (GSM4732270, GSM4732271, GSM4732272) after GSI washout was used, and merged PRO-seq data from all conditions in this work were used for this analysis, to comprehensively capture gene activity in these samples.

### Differential gene expression analysis

TT-seq reads mapping to exons in the active gene models were counted using the featurecounts function in the Rsubread^50^ package version 2.14.2, and these values were normalized using size factors in DESeq2^47^ version 1.40.2. Significance values were calculated for each comparison between conditions using the design formula ∼replicate + condition. Genes were called significantly changed if they had adjusted p value < 0.01 and |FC| > 1.5 in either the 1h_washout_DMSO or the 4h_washout_DMSO condition vs 0h_GSI_DMSO. 4 groups were then assigned: *<u>Notch upregulated</u>* (significantly upregulated in 1h_washout_DMSO vs 0h_GSI_DMSO and (1h_washout_DMSO / 1h_GSI_DMSO) > 1.5 OR significantly upregulated in 4h_washout_DMSO vs 0h_GSI_DMSO and (4h_washout_DMSO / 4h_GSI_DMSO) > 1.5 ; *<u>Notch</u> <u>downregulated</u>* (significantly downregulated in 1h_washout_DMSO vs 0h_GSI_DMSO and (1h_washout_DMSO / 1h_GSI_DMSO) < 0.667 OR significantly upregulated in 4h_washout_DMSO vs 0h_GSI_DMSO and (4h_washout_DMSO / 4h_GSI_DMSO) < .667; *<u>Nonspecific upregulated</u>* (significantly upregulated in 1h_washout_DMSO vs 0h_GSI_DMSO and not *Notch upregulated*; *<u>Nonspecific downregulated</u>* (significantly upregulated in 1h_washout_DMSO vs 0h_GSI_DMSO and not *Notch downregulated*. Genes were then filtered to include only those with at least 60 PRO-seq promoter (TSS to TSS+150) reads in one condition, to select only for high confidence genes. For display in heatmaps, experimental replicates were averaged, and gene expression was normalized such that the highest condition being displayed was set to 1.

A list of unchanged genes was defined as genes not significant in any of the above four categories that additionally had a |FC| < 1.10 in either the 1h_washout_DMSO or the 4h_washout_DMSO condition vs 0h_GSI_DMSO (n=1484).

A list of inactive genes was defined as genes with fewer than 10 PRO-seq reads within the promoter (TSS to TSS + 150 nt) at any time point (n=1295).

### Enhancer identification

PRO-seq single-nucleotide bedgraph files were merged over all time points, and converted into bigwig files. dREG^51^ was used to predict putative regulatory elements, using these input files. These were refined using a custom filtering script, dRIP-filter, to keep only peaks with a dREG score of at least 0.5, a p-value of less than 0.025, and at least 5 PRO-seq reads in 2 conditions. As an additional validation of enhancer quality, H3K27ac ChIP signal (merged over all time points) was counted over the potential dREG peaks using makeheatmap using the “-b v -v t -s b” options, and only peaks with at least 300 reads were kept. Next, promoter-proximal peaks (within 1kb of active annotated promoters) were removed, leaving only distal enhancers, and enhancers within 1kb of each other were merged into one window using bedtools merge. Enhancers overlapping rRNA, snRNAs, scRNAs, srpRNAs, tRNAs, or snoRNAs were removed using bedtools intersect. Enhancers were then classified as intragenic if they overlap an active gene, or intergenic if they did not. Intragenic enhancers overlapping two genes on opposite strands were discarded, leaving a final list of 12,040 intergenic and 8,983 intragenic enhancers.

### Notch transcription complex (NTC) binding analysis

8,533 ChIP peaks bound by both RBPJ and MAML1, as determined by Pan et al.^17^, were defined as NTC binding regions. The intersectBed tool was used to identify genes with promoter NTC binding, those with an NTC binding region overlapping the 1kb upstream of the TSS. To identify NTC-bound enhancers, intersectBed was used to label enhancers that overlap an NTC binding region. The distance from a gene TSS to the closest enhancer was calculated using the bedtools closest function.

### Assigning enhancers to genes

The closest NTC-bound enhancer (see above) to the TSS of each Notch-upregulated gene was identified using the bedtools closest function. 3 genes had no NTC-bound enhancers within 1,000,000 bp of the TSS, and were excluded from further analysis, yielding a group of 58 closest enhancers to Notch-upregulated genes, shown in Figures 3G-H. For aggregate plots shown in 3E-F, the dominant enhancer TSS for each of these 58 enhancers was identified in order to center the plots. Unannotated TSSs called by TSS-call as part of the genome annotation pipeline (see above) within the dREG peaks were identified using bedtools intersect, and the enhancer with the highest TSS score (from TSS-call) for each enhancer was selected as the dominant TSS. For intragenic enhancers, only TSSs on the non-gene strand were considered. This resulted in a list of 56 enhancers with a dominant TSS, which are plotted in Figures 3E-F.

### Gene Ontology Enrichment

Biological Process GO Term enrichment was determined using the Molecular Signatures Database (MSigDB)^52^ website (www.gsea-msigdb.org), on all genes in Groups 1 and 2. For groups 3 and 4, which had more than 500 genes, the top 500 genes with largest (Group 3) or smallest (Group 4) fold change at 4 hours were used as the input.

### Summing reads over genomic features

For analysis of PRO-seq reads over genes, genomic windows were defined starting from the active gene annotation. Gene body regions were defined as TSS + 250 nt to TSS + 5 kb or the first intragenic enhancer, whichever came first. Principal Component Analysis shown in Figure S2 was performed over these regions for genes over 1 kb in length. TSS windows were defined as TSS to TSS + 150 nt. Reads were counted over these regions using the custom script makeheatmap using the “-b v -l s -s s” options. Counts were normalized by the length of the genomic window.

Total ATAC-seq reads at promoters were counted using makeheatmap using the “-b v -v t -s b” options, over a window beginning 450 bp upstream of the TSS to 149 nt downstream of the TSS. This window was chosen the focus on the nucleosome depleted region at the promoter.

Total PRO-seq reads over dREG peaks for figure 3H were counted using makeheatmap using the “-b v -l s -s b” option for intergenic peaks, and “-b v -l s -s o” for intragenic peaks to avoid conflating gene reads with the enhancer reads. H3K27ac ChIP-seq and ATAC-reads reads over dREG peaks for figures 3I and 3J were counted using makeheatmap using the “-b v -v t -s b” options on single-nucleotide bedgraph files merged over replicates. The full dREG peak region was chosen for this analysis.

### Aggregate plots

For PRO-seq aggregate plots around promoters, PRO-seq reads were counted in 20 nt bins from TSS −100 nt to TSS + 1 kb, summing the total reads within the window for each gene using makeheatmap with the “-b c -a s -l s -v t -s s” options. For aggregate plots in the gene body, PRO-seq reads were counted in 40 nt bins from TSS + 500 nt to TSS + 1 kb, summing the total reads within the window for each gene. The average value over the Notch upregulated genes for each bin is shown in the plots.

For ATAC-seq aggregate plots at promoters, ATAC-seq reads were counted in 75bp bins within 1 kb of the gene TSS using makeheatmap with the “-b c -a u -v t -s b” options.

For ATAC-seq and ChIP-seq aggregate plots over dREG peaks, reads were counted around the dominant TSS for the enhancer using makeheatmap with the “-b c -a u -v t -s b” options. ATAC-seq reads were counted in 75 bp bins, and ChIP-seq reads were counted in 50 bp bins.

For PRO-seq aggregate plots over dREG peaks, PRO-seq reads were counted in 75 nt bins around the dominant TSS for the enhancer using makeheatmap with “-l s -b c -a u -v t -s s” options for intragenic enhancers (to only count reads on the strand with the TSS, the opposite strand of the genic reads), or “-l s -b c -a u -v t -s b” options for the intergenic enhancers. The reads per bin for the intergenic enhancers were divided by 2 to estimate the reads per strand to enable the averaging of reads over all enhancers. The average signal per bin over the 56 enhancers with TSSs are plotted.

### Calculating pausing indices

Pausing index is defined as PRO-seq reads per base pair in the TSS (TSS to TSS + 150 nt) divided by PRO-seq reads per base pair in TSS + 250 nt to TSS + 2250 nt or TES, whichever comes first. Reads in these regions were counted using makeheatmap using the “-b v -l s -s s” options.

## ACKNOWLEDGMENTS

We thank Marie Bao and members of the Blacklow and Adelman labs for helpful discussion. We thank the Nascent Transcriptomics Core at Harvard Medical School for performing PRO-seq and TT-seq library construction, Apoorva Baluapuri, the Harvard Medical School Biopolymers Facility, and the Bauer Core Facility at Harvard University for sequencing.

This work was supported by NIH award R35 CA220340 to S.C.B., R01GM139960 to K.A., a National Science Foundation Graduate Research Fellowship (DGE1745303) and the Sophia H.Y Chang Fellowship to C.A.M., a CIHR Banting Postdoctoral Fellowship to B.J.E.M., and a LLS Career Development Fellowship and K99GM144750 grant to J.M.R.

## AUTHOR CONTRIBUTIONS

Conceptualization, Methodology, S.C.B., K.A., J.M.R, J.C.A; Investigation, Data Curation, and Visualization, J.M.R.; Data Analysis, J.M.R., C.A.M., B.J.E.M; Resources, A.P.M.; Writing. J.M.R., S.C.B., K.A.; Supervision, K.A., S.C.B., Funding Acquisition, J.M.R., S.C.B

## SUPPLEMENTARY METHODS

### Keratinocyte Media Composition

Media for SC2 culture consisted of: 3:1 DMEM:F12 + 10% Fetal Bovine Serum (Gemini Bio-Products 100-106) + 1X Penicillin-Streptomycin (ThermoFisher 15140163) + 1X Reagent Mix (see below) + 1 μM Compound E (EMD Millipore 565790) gamma secretase inhibitor (GSI). 100X Reagent Mix was prepared in 3:1 DMEM:F12 media, containing 40 μg/mL Hydrocortisone (Sigma-Aldrich H0888), 500 ng/mL Insulin (Sigma-Aldrich I6634), 1μg/mL EGF (Sigma-Aldrich E9644), 0.84 μg/μL Cholera toxin (Sigma-Aldrich C9903), 500 μg/mL Transferrin (Sigma-Aldrich T2036), and 1.3 μg/mL Liothyronine (Sigma-Aldrich T6397).

### RNA purification for TT-seq library construction

Fly (S2) cells used for spike-in were labeled for 2 hr with 4-SU (500 µM), and harvested in Trizol. 5 % 4-SU labeled spike-in cells (e.g. 0.5 million fly cells into 10 million human cells) were added directly to the Trizol lysate for each experimental condition. This was then split into two 700 µL tubes for purification. First the lysate was homogenized using a QiaShredder column (Qiagen 79656). 150 µL of chloroform was added to the homogenate, which was then vortexed, incubated for 5 minutes, and spun down at 12,000 *g* for 15 minutes at 4°C to separate the phases. 360 μL of the aqueous phase was transferred to a new tube, and 540 µL of ethanol with 1.5 mM DTT was added. This material was then applied to miRNeasy mini kit column (Qiagen 217004), and RNA was isolated following the kit protocol, using an on-column digestion with DNase (Qiagen 79254). After the isolation, the two columns per sample were re-combined. Aliquots of RNA were removed for quantification by spectrophotometry and analysis of RNA integrity by Agilent TapeStation 4200 using RNA high sensitivity tapes. All samples were confirmed to have RNA integrity numbers (RIN) > 9.0 before proceeding to TT-seq library construction.

### TT-seq library construction

50 µg of RNA per sample was brought to a volume of 80 µL with nuclease-free water and placed on ice. RNA was then lightly fragmented by addition of 20 µL cold 5X fragmentation solution (375 mM Tris-HCl (pH 8.3), 562.5 mM KCl, 22.5 mM MgCl_2_) and incubation at 94°C for 2 minutes 15 seconds. At the end of the fragmentation time, RNA was placed immediately on ice and 25 µL of cold 250 mM EDTA was added. RNA was precipitated by addition of 1/10 volume of 5 M NaCl, 2.5 volumes of 100 % ethanol and incubation at −20°C overnight. RNA was pelleted, washed, quantified, and analyzed again as above. Fragmented RNA was biotinylated essentially as described in Duffy et al.^53^ with the following modifications: the biotinylation reaction was performed in a total volume of 200 µL and allowed to incubate for 45 min in the dark. Excess biotin was removed using chloroform:isoamyl alcohol as per Dölken et al.^54^ and MaXtract tubes (Qiagen 129056) were used to separate organic and aqueous phases. Biotinylated RNA was resuspended in 100 µL of nuclease-free water and aliquots taken to use as the total RNA input fraction. In parallel, Dynabeads M-280 Streptavidin (ThermoFisher 11205D) were prepared for binding to render them RNase-free: for each sample, 75 µL of beads were used and treated in batch to render them RNase free. The beads were incubated 10min in a solution of 100 mM NaOH, 50 mM NaCl, placed on a magnetic stand, and then washed, resuspending the beads fully for each wash, twice with 500 µL of 100 mM NaCl, twice with 1 X TT-seq wash solution (100 mM Tris-HCl (pH 7.5), 10mM EDTA, 1 M NaCl, 0.05% Tween 20 in nuclease free water to which 1 µL SUPERase-In RNase Inhibitor (ThermoFisher AM2696) per 5mL solution is added prior to use), once in 0.3 X TT-seq wash solution, and finally resuspended in 52 µL/sample of 0.3 X TT-seq wash solution + 1 µL/sample SuperaseIn RNase Inhibitor.

Biotinylated RNA was heated at 65°C for 5min, placed on ice for 2 min, and mixed with 50 µL of prepared beads. Samples were rotated at room temperature in the dark for 30 min. After binding, the tubes were placed on a magnetic rack and the beads were washed 4 times with 500 µL of 1 X TT-seq wash solution to remove unbound RNA, fully resuspending for each wash. The wash solution was removed, and the beads resuspended in 50 µL of 0.1 M DTT (freshly diluted from 1 M stock) and rolled in the dark for 15 min at room temp. The eluted RNA was recovered and the elution step repeated with an additional 50 µL of 0.1 M DTT. The combined eluates were purified using the RNA clean-up and concentration microElute kit (Norgen. 61000) following the manufacturer’s instructions for small RNA enriched samples. Final elution was performed in 14 µL of nuclease-free water and the eluate was reapplied to the column for a total of 2 elution steps. A Qubit RNA high sensitivity reagent kit was used to quantify the input RNA and enriched RNA. Yields of 1-2 % were typical. 150 ng of enriched RNA was used for library construction with the NEBNext Ultra II Directional RNAseq kit (New England Biolabs E7765S), with the rRNA depletion module (New England Biolabs E7405L), and unique dual index set 2 (New England Biolabs E6442S). Libraries were pooled and sequenced paired-end with the 200 cycle kit on a Novaseq S4 single lane.

### ChIP-seq chromatin preparation

To crosslink cells, at the appropriate time after washout, media was removed from the plates and 10 mL PBS was added (PBS contained 1 μM GSI for the mock washout sample). 9 mL of 2.1 % formaldehyde in PBS was added to the plate (1% final concentration) (3 plates / sample were staggered by 15 seconds to ensure equal crosslinking time), and cells were fixed on a shaker for precisely 15 min. 1 mL 2.5 M Glycine was added after 15 min, plates were incubated for an additional 5 min, and then placed on ice. Cells were scraped off the plate and collected in a 50 mL falcon tube. Plates were rinsed with 20 mL cold PBS, which was added to the falcon tube. Cells were pelleted at 300 *g* for 5 min at 4°C, supernatant was discarded. The 3 tubes/sample were combined and resuspended in 40 mL of PBS, pelleted at 300 *g* for 5 min at 4°C, resuspended in 10 mL PBS, and pelleted again. The pellet was then resuspended in 250 μL of sonication buffer (20 mM Tris (pH 8.0), 2 mM EDTA, 0.5 mM EGTA, 0.5 % SDS, 0.5 mM PMSF, and 1 mini protease inhibitor tablet per 10 mL buffer. Chromatin was incubated on ice for 10min, 2 5 μL samples were taken for chromatin normalization, and then samples were flash frozen and stored at −80°C.

To normalize chromatin, the 5 μL sample was added to 200 μL QuickExtract Buffer (VWR). Samples were vortexed 15 s, heated at 65°C at 1500 rpm for 15 min, vortexed again for 15 s, then heated at 98°C for 6 minutes. A standard sample of SC2 cells at 10^8^ cells/mL was processed in this same way to make a standard. The DNA concentration of all samples and the standards were measured by the Qubit dsDNA high sensitivity quantitation kit (ThermoFisher), and samples were adjusted to the 10^8^ cells/mL standard with fresh sonication buffer.

Chromatin was then sonicated using a QSonica sonicator (70 % amplitude, 15 seconds ON / 45 seconds off cycles) to an average fragment size of ∼200 bp, flash frozen, and stored at - 80°C. Chromatin from 2.5 million cells was diluted in 1 mL IP buffer (20 mM Tris (pH 8.0), 2 mM EDTA, 0.5% Triton X-100, 150 mM NaCl, 10% Glycerol), and precleared for 2 hours at 4°C, rotating with 30 μL Protein A agarose beads (Millipore Sigma 16-125) that had been pre-equilibrated in cold IP buffer. Precleared input was transferred to a new tube, 250 μL additional IP buffer and 30 μL H3K27ac antibody (Active Motive 39133) were added, and samples were incubated overnight at 4°C, rotating. 200 μL Protein A beads were added to each IP, and samples were incubated for 2 hours at 4°C, rotating. Samples were washed once with Low-Salt Buffer (20 mM Tris (pH 8.0), 2 mM EDTA, 1% Triton X-100, 150 mM NaCl, 0.1% SDS), three times with High-Salt Buffer (20 mM Tris pH 8.0, 2 mM EDTA, 1% Triton X-100, 500 mM NaCl, 0.1% SDS), once with Lithium Chloride Buffer (20 mM Tris pH 8.0, 2 mM EDTA, 250 mM LiCl, 1% IGEPAL CA-630, 1% sodium deoxycholate), and twice with TE Buffer (10 mM Tris-HCl, 1 mM EDTA). For each wash, samples were rotated for 3 minutes with 1 mL of cold wash solution. Samples were eluted twice by rotating beads for 15 min at room temperature with 250 μL elution buffer (1 % SDS, 100 mM NaHCO_3_). The two elutions were combined, NaCl was added to 200 mM, and samples were incubated overnight at 65°C to reverse crosslinks. Samples were treated with Proteinase K (New England Biolabs P8107) for 30 min, extracted with phenol:chloroform:isoamyl alcohol, and resuspended in 100 μL water.

### ATAC-seq sample preparation and library construction

ATAC was performed according to the OmniATAC protocol^44^. Briefly, cells were pelleted at 500 *g* for 5 min at 4°C, and supernatant was removed. The cell pellet was resuspended in 50 μL ATAC lysis buffer by pipetting up and down 3 times (Cold ATAC-Resuspension Buffer (10 mM Tris–HCl pH 7.5, 10 mM NaCl and 3 mM MgCl2) + 0.1 % NP40 (EMD Millipore 492018) + 0.1 % Tween-20 (EMD Millipore 655206) + 0.01 % digitonin (Promega G9441)). Up to 4 samples were processed at 1 time, and samples were staggered by 15 seconds. After exactly 3 min of lysis, 1 mL of ATAC wash buffer (Cold ATAC-Resuspension Buffer + 0.1 % Tween-20 (EMD Millipore 655206)) was added to the tube. Nuclei were pelleted at 500 *g* for 10 min at 4°C, and supernatant was removed. Nuclei were resuspended in 50 μL of Transposition mix (1x TD buffer (Illumina 20334197), 3 μL Tn5 transposase (Illumina 20334197), 0.33x PBS, 0.01 % digitonin, 0.1 % Tween-20), and incubated at 37°C in a thermomixer at 1000 rpm for 30 minutes. The reactions were quenched with 250 μL of binding buffer from the MinElute PCR purification kit (Qiagen 28004) and purified using this kit. This was then used as the input for barcoding PCR, to add sequencing adapters and sequencing barcodes, as described in Grandi et al.^44^. Both the qPCR and Qubit methods were used to quantify the libraries to determine the total number of PCR cycles, as described in the OmniATAC protocol. After the final PCR cycles, libraries were cleaned up with the MinElute PCR purification kit, quantified using the NEBNext Library Quant kit (New England Biolabs E7630), and pooled for sequencing.

### PRO-seq cell permeabilization

Cells were pelleted at 300 *g* for 4 min at 4°C, supernatant was removed, and cells were resuspended in 10 mL cold PBS. Again, cells were pelleted at 300 *g* for 4 min at 4°C, supernatant was removed, and cells were resuspended in 250 μL cold Buffer W (10 mM Tris-Cl, pH 8.0, 10 mM KCl, 250 mM sucrose, 5 mM MgCl_2_, 1 mM EGTA, 10% (w/v) glycerol in DEPC ddH_2_O + 1 protease inhibitor tablet (Millipore Sigma 5056489001), 0.5 mM DTT, and 10 μL SUPERase-In RNase inhibitor per 50 mL buffer added fresh before beginning the experiment). 10 mL of Buffer P (10 mM Tris-Cl, pH 8.0, 10 mM KCl, 250 mM sucrose, 5 mM MgCl_2_, 1 mM EGTA, 0.05 % Tween-20 10 % (w/v) glycerol in DEPC ddH_2_O + 1 protease inhibitor tablet, 0.5 mM DTT, and 10 μL SUPERase-In RNase inhibitor per 50 mL buffer added fresh before beginning the experiment) was slowly added to the tube, and cells were incubated for 5 minutes on ice to permeabilize. Two times, cells were pelleted at 400 *g* for 4 min at 4°C, supernatant was removed, and cells were resuspended in 1 mL Buffer W, and pipetted up and down slowly with a wide-bore pipette tip. An additional 9 mL of Buffer W was added. After the second wash, cells were pelleted at 500 *g* for 4 min at 4°C, resuspended in 250 μL freezing buffer (10 mM Tris-Cl pH 8, 40% (w/v) glycerol, 5 mM MgCl_2_, 1.1 mM EDTA + 1 protease inhibitor tablet, 0.5 mM DTT, and 10 μL SUPERase-In RNase inhibitor per 50 mL buffer added fresh before beginning the experiment), and moved to a 1.5 mL tube. The 15 mL tube was rinsed with another 250 μL freezing buffer, which was combined to make 500 μL of permeabilized cells in freezing buffer, which were snap frozen in liquid nitrogen and stored at −80°C.

### PRO-seq library construction

PRO-seq library construction was performed by the Nascent Transcriptomics Core at Harvard Medical School, Boston, MA. Aliquots of frozen permeabilized cells were thawed on ice and pipetted gently to fully resuspend. Aliquots were removed and permeabilized cells were counted using a Luna II, Logos Biosystems instrument. For each sample, 1 million permeabilized cells were used for nuclear run-on, with 50,000 permeabilized *Drosophila* S2 cells added to each sample for normalization. Nuclear run-on assays and library preparation were performed essentially as described in Reimer et al.^55^ with modifications noted: 2X nuclear run-on buffer consisted of (10 mM Tris (pH 8), 10 mM MgCl2, 1 mM DTT, 300 mM KCl, 20 μM/each biotin-11-NTPs (Perkin Elmer NEL54(2/3/4/5)001), 0.8 U/μL SuperaseIN, 1 % sarkosyl (Sigma-Aldrich L7414). Run-on reactions were performed at 37°C. Fragmentation of RNA was performed in 100 mM NaOH at 0°C for 10 minutes and stopped by addition of one volume of ice cold 1 M Tris (pH 6.8). Adenylated 3’ adapter was prepared using the 5’ DNA adenylation kit (New England Biolabs E2610S) and ligated using T4 RNA ligase 2, truncated KQ (NEB, per manufacturer’s instructions with 15 % PEG-8000 final) and incubated at 16°C overnight. 180 μL of betaine blocking buffer (1.42 g of betaine brought to 10 mL with binding buffer supplemented to 0.6 μM blocking oligo (TCCGACGATCCCACGTTCCCGTGG/3InvdT/)) was mixed with ligations and incubated 5 min at 65°C and 2 min on ice prior to addition of Dynabead M-280 streptavidin beads (ThermoFisher 11205D). After T4 polynucleotide kinase (New England Biolabs M0201) treatment, beads were washed once each with high salt, low salt, and blocking oligo wash (0.25X T4 RNA ligase buffer (New England Biolabs M0437), 0.3 μM blocking oligo) solutions and resuspended in 5’ adapter mix (10 pmol 5’ adapter, 30 pmol blocking oligo, water). 5’ adapter ligation was per Reimer et al.^55^, but with 15 % PEG-8000 final. Eluted cDNA was amplified 5-cycles (NEBNext Ultra II Q5 master mix (New England Biolabs M0544) with Illumina TruSeq PCR primers RP-1 and RPI-X) following the manufacturer’s suggested cycling protocol for library construction. A portion of preCR was serially diluted and for test amplification to determine optimal amplification of final libraries. Pooled libraries were sequenced using the Illumina 200 cycle kit on an S4 single lane, followed by additional run on a full SP lane for more depth for some samples.

### TT-seq mapping

Read pairs were trimmed to 100 base pairs using a custom script (trim_and_filter_PE.pl), and reads with a minimum average quality score of 20 were kept. Reads were first mapped to the spike genome (dm6) using bowtie1.2.2 (-n2 -l 40 -X1000 -p 5 --best -3 1 parameters). Reads not mapping to the spike genome were mapped to the human genome (hg38) using with STAR2.7.3a^47^, using parameters --quantMode TranscriptomeSAM GeneCounts --outSAMtype BAM SortedByCoordinate --limitBAMsortRAM 42949672960 --outMultimapperOrder Random -- outSAMattrIHstart 0 --outFilterType BySJout --outFilterMismatchNmax 4 --alignSJoverhangMin 8 --outSAMstrandField intronMotif --outFilterIntronMotifs RemoveNoncanonicalUnannotated -- alignIntronMin 20 --alignIntronMa× 1000000 --alignMatesGapMa× 1000000 --outWigType bedGraph --outWigNorm None --outFilterScoreMinOverLread 0 –outFilterMatchNminOverLread 0. Duplicates were also removed using STAR, and stranded coverage bedGraph files were generated from deduplicated BAM files using STAR.

To normalize samples, reads mapping to exons in the active gene models (see below) were counted using featurecounts, and used to calculate size factors using DESeq2. PCA plots were generated using DESeq2 over the top 500 genes. These size factors were scaled so the minimum was 1, and used to normalize bedGraphs with the custom script normalize_bedGraph.pl. Biological replicates (n=2) were merged using the script bedgraphs2stbedGraph.pl.

| Sample Name | Total | Reads mapping to hg38 | Reads mapping to dm6 | Size Factor | Normalization Factor |
| --- | --- | --- | --- | --- | --- |
| 0h_GSI_BRM014_rep1 | 150088510 | 138301345 | 150088510 | 1.05 | 1.22 |
| 0h_GSI_BRM014_rep2 | 129512568 | 118460711 | 13401548 | 1.05 | 1.22 |
| 0h_GSI_DMSO_rep1 | 113814982 | 103229584 | 16067730 | 0.88 | 1.03 |
| 0h_GSI_DMSO_rep2 | 127092978 | 117016818 | 10177307 | 0.94 | 1.10 |
| 1h_GSI_BRM014_rep1 | 138189730 | 124468545 | 25550404 | 1.07 | 1.25 |
| 1h_GSI_BRM014_rep2 | 153527552 | 142323388 | 8963298 | 1.10 | 1.28 |
| 1h_GSI_DMSO_rep1 | 124021166 | 112513927 | 18322070 | 1.01 | 1.18 |
| 1h_GSI_DMSO_rep2 | 126653138 | 117185129 | 7514158 | 0.94 | 1.10 |
| 1h_washout_BRM014_rep1 | 149533912 | 136881840 | 18589668 | 1.13 | 1.32 |
| 1h_washout_BRM014_rep2 | 126805849 | 116166177 | 8990111 | 0.90 | 1.05 |
| 1h_washout_DMSO_rep1 | 206317803 | 191540602 | 13777718 | 1.38 | 1.61 |
| 1h_washout_DMSO_rep2 | 141587149 | 130485870 | 10945308 | 1.03 | 1.20 |
| 4h_GSI_BRM014_rep1 | 113177888 | 103802452 | 10233517 | 0.86 | 1.00 |
| 4h_GSI_BRM014_rep2 | 139996078 | 129243376 | 7944409 | 1.02 | 1.19 |
| 4h_GSI_DMSO_rep1 | 137548171 | 125334006 | 15977982 | 1.03 | 1.21 |
| 4h_GSI_DMSO_rep2 | 125876159 | 114811952 | 9318722 | 0.95 | 1.11 |
| 4h_washout_BRM014_rep1 | 114345951 | 104417618 | 13314893 | 0.90 | 1.05 |
| 4h_washout_BRM014_rep2 | 138655109 | 127374492 | 9022958 | 1.07 | 1.25 |
| 4h_washout_DMSO_rep1 | 127331045 | 117044027 | 10629014 | 0.97 | 1.13 |
| 4h_washout_DMSO_rep2 | 121877894 | 111587627 | 8130586 | 0.94 | 1.09 |

### ChIP-seq mapping

Read 1 was trimmed to positions 2-49 using a custom script (trim_and_filter_SE.pl), and reads with a minimum average quality score of 20 were kept. ChIP-seq mapping statistics are indicated below.

| Sample Name | Total | Reads mapping to hg38 | Reads mapping to dm6 | Normalization Factor |
| --- | --- | --- | --- | --- |
| 0h_repA | 86958599 | 77821951 | 150676 | 1.33 |
| 0h_repB | 84908653 | 76828968 | 154255 | 1.31 |
| 0h_input_repA | 84047641 | 75998979 | 119330 | 1.33 |
| 0h_input_repB | 79677737 | 73910713 | 67541 | 1.32 |
| 15m_repA | 85386928 | 74785415 | 170064 | 1.27 |
| 15m_repB | 81909319 | 73974099 | 85509 | 1.27 |
| 30m_repA | 74940561 | 66308039 | 188874 | 1.12 |
| 30m_repB | 92092844 | 81888573 | 138717 | 1.37 |
| 45m_repA | 83365374 | 73053116 | 150563 | 1.22 |
| 45m_repB | 91860316 | 81491941 | 128015 | 1.37 |
| 1h_repA | 87400524 | 76671688 | 180077 | 1.28 |
| 1h_repB | 131159821 | 115050788 | 506225 | 1.69 |
| 90m_repA | 66913671 | 59101893 | 138359 | 1.00 |
| 90m_repB | 86418798 | 75985450 | 282378 | 1.20 |
| 2h_repA | 67841126 | 60113642 | 171034 | 1.02 |
| 2h_repB | 93271276 | 81119922 | 141842 | 1.33 |
| 4h_repA | 91720189 | 80490390 | 194821 | 1.27 |
| 4h_repB | 101779012 | 89968296 | 111693 | 1.44 |
| 4h_input_repA | 96217572 | 88365662 | 133355 | 1.51 |
| 4h_input_repB | 80521699 | 72713558 | 65706 | 1.28 |

### ATAC-seq mapping

Read pairs were trimmed to 100 base pairs using a custom script (trim_and_filter_PE.pl), and reads with a minimum average quality score of 20 were kept. Adapter sequences were removed using cutadapt 1.14^56^. Reads were first mapped to the spike genome (dm6), using bowtie1.2.2 (-k1 -v2 -X1000 – allow-contain -- best parameters). Reads that did not map to the spike genome were mapped to the human genome (hg38), using bowtie1.2.2^42^ (-k1 -v2 -X1000 – allow-contain -- best parameters). Samtools 1.9^57^ was used to flag duplicate reads, which were then removed. A custom script, extract_fragments.pl, was used to filter and retain unique reads between 10 and 150 base pairs, which correspond to regions of open chromatin, and convert files into bedGraph format. Spike return rates were not significantly different between samples. To normalize samples, reads mapping to the −1 kb to +1 kb region around active promoters in this cell type (see Genome Annotation below) were counted using the custom script makeheatmap with the “-b v -v t -s b” options. The number of these reads was used to calculate size factors using DESeq2^47^. PCA plots were generated using DESeq2 over the top 500 promoters. These size factors were scaled so the minimum was 1, and used to normalize bedGraphs with the custom script normalize_bedGraph.pl. Biological replicates (n=2) were merged using the script bedgraphs2stbedGraph.pl.

| Sample Name | Total | Reads mapping to hg38 | Reads mapping to dm6 | Reads within fragment size cutoff | Size Factor | Normalization Factor |
| --- | --- | --- | --- | --- | --- | --- |
| 0h_GSI_BRM014_rep1 | 34202112 | 31976690 | 227554 | 8708132 | 1.39 | 5.09 |
| 0h_GSI_BRM014_rep2 | 33906592 | 31638459 | 82304 | 8096329 | 1.40 | 5.15 |
| 0h_GSI_DMSO_rep1 | 29207488 | 27429824 | 76822 | 8036150 | 0.79 | 2.90 |
| 0h_GSI_DMSO_rep2 | 36418596 | 33858170 | 153535 | 8362786 | 1.81 | 6.63 |
| 1h_GSI_BRM014_rep1 | 28689136 | 26775117 | 103456 | 5375668 | 1.02 | 3.73 |
| 1h_GSI_BRM014_rep2 | 39727072 | 36959808 | 36893 | 9657709 | 0.97 | 3.53 |
| 1h_GSI_DMSO_rep1 | 34142160 | 32125674 | 7737 | 9365400 | 0.77 | 2.81 |
| 1h_GSI_DMSO_rep2 | 35218200 | 33117630 | 30062 | 7867053 | 1.69 | 6.20 |
| 1h_washout_BRM014_rep1 | 33987051 | 31724447 | 60562 | 9456331 | 0.87 | 3.20 |
| 1h_washout_BRM014_rep2 | 42124438 | 39171424 | 31658 | 9655117 | 1.48 | 5.43 |
| 1h_washout_DMSO_rep1 | 17165045 | 15752974 | 42975 | 2300391 | 0.27 | 1.00 |
| 1h_washout_DMSO_rep2 | 38410233 | 35726823 | 82631 | 9985688 | 0.77 | 2.82 |
| 2h_GSI_BRM014_rep1 | 31547139 | 29232600 | 136633 | 7320975 | 1.13 | 4.12 |
| 2h_GSI_BRM014_rep2 | 30714106 | 28227978 | 101475 | 6101677 | 1.44 | 5.27 |
| 2h_GSI_DMSO_rep1 | 33833327 | 31810936 | 111365 | 8569085 | 1.44 | 5.28 |
| 2h_GSI_DMSO_rep2 | 26264980 | 24429806 | 81237 | 6478067 | 1.02 | 3.72 |
| 2h_washout_BRM014_rep1 | 33069311 | 30559738 | 129552 | 6115721 | 1.57 | 5.75 |
| 2h_washout_BRM014_rep2 | 32665148 | 30221771 | 14722 | 7050846 | 1.28 | 4.66 |
| 2h_washout_DMSO_rep1 | 32355097 | 30206171 | 36064 | 8557728 | 0.67 | 2.44 |
| 2h washout DMSO_rep2 | 26417570 | 24386901 | 21189 | 6835309 | 0.62 | 2.25 |
| 4h GSI BRM014_rep1 | 36971274 | 34566802 | 60087 | 10780739 | 0.95 | 3.46 |
| 4h GSI BRM014_rep2 | 30014033 | 28110552 | 19452 | 7502842 | 0.99 | 3.62 |
| 4h GSI DMSO_rep1 | 39905542 | 37480566 | 29108 | 10964823 | 0.82 | 3.01 |
| 4h GSI DMSO_rep2 | 34191134 | 32284375 | 15791 | 8784192 | 1.25 | 4.57 |
| 4h washout BRM014_rep1 | 34567268 | 31890198 | 117786 | 9651176 | 1.24 | 4.54 |
| 4h washout BRM014_rep2 | 31650134 | 29445166 | 22535 | 7502925 | 0.98 | 3.57 |
| 4h washout DMSO_rep1 | 29269199 | 27271393 | 12857 | 9395100 | 0.58 | 2.04 |
| 4h washout DMSO_rep2 | 32502539 | 30536084 | 48340 | 9485869 | 1.25 | 4.59 |

### PRO-seq mapping

Read pairs were trimmed using cutadapt 1.14^56^ to remove adapter sequences (-O 1 -- match-read-wildcards -m {20}). An additional nucleotide was removed from the end of read 1 (R1), using seqtk trimfq (https://github.com/lh3/seqtk), to preserve a single mate orientation during alignment. The paired end reads were then mapped to a combined genome index, including both the spike (dm6) and primary (hg38) genomes, using bowtie2^48^. Properly paired reads were retained. These read pairs were then separated based on the genome (i.e. spike-in vs primary) to which they mapped. Reads mapping to the reference genome were separated according to whether they were R1 or R2, sorted via samtools^57^ 1.3.1 (-n), and subsequently converted to bedGraph format using a custom script (bowtie2stdBedGraph.pl). We note that this script counts each read once at the exact 3’ end of the nascent RNA. Because R1 in PRO-seq reveals the position of the RNA 3’ end, the “+” and “-” strands were swapped to generate bedGraphs representing 3’ end positions at single nucleotide resolution.

| Sample Name | Total | Reads mapping to hg38 | Reads mapping to dm6 | Normalization Factor |
| --- | --- | --- | --- | --- |
| 0h_rep1 | 85970282 | 76152127 | 2040865 | 1.54 |
| 0h_rep2 | 103531128 | 93818174 | 2346439 | 1.90 |
| 0h_rep3 | 88225365 | 75932963 | 2660924 | 1.54 |
| 15min_rep2 | 57356757 | 49456531 | 2394454 | 1.00 |
| 15min_rep3 | 93557006 | 83885924 | 2891559 | 1.70 |
| 30min_rep1 | 77477601 | 68196399 | 3208443 | 1.38 |
| 30min_rep2 | 78302067 | 67738226 | 2573842 | 1.37 |
| 45min_rep1 | 76402172 | 69977167 | 1972974 | 1.41 |
| 45min_rep2 | 88623510 | 79516628 | 1802120 | 1.61 |
| 1h_rep1 | 101092654 | 91020976 | 1955255 | 1.84 |
| 1h_rep2 | 101437843 | 93250008 | 2100768 | 1.89 |
| 90min_rep1 | 101175352 | 92626064 | 2283885 | 1.87 |
| 90min_rep2 | 109737877 | 99051682 | 2718587 | 2.00 |
| 2h_rep1 | 95528400 | 86606556 | 1974717 | 1.75 |
| 2h_rep2 | 88938370 | 80741577 | 1760960 | 1.63 |
| 3h_rep1 | 90661683 | 82416142 | 2035634 | 1.67 |
| 3h_rep2 | 124589832 | 113193709 | 2539083 | 2.29 |
| 4h_rep1 | 90560540 | 82816019 | 1332947 | 2.03 |
| 4h_rep2 | 109462234 | 100521172 | 2378508 | 1.67 |

---

**Figure S1.**
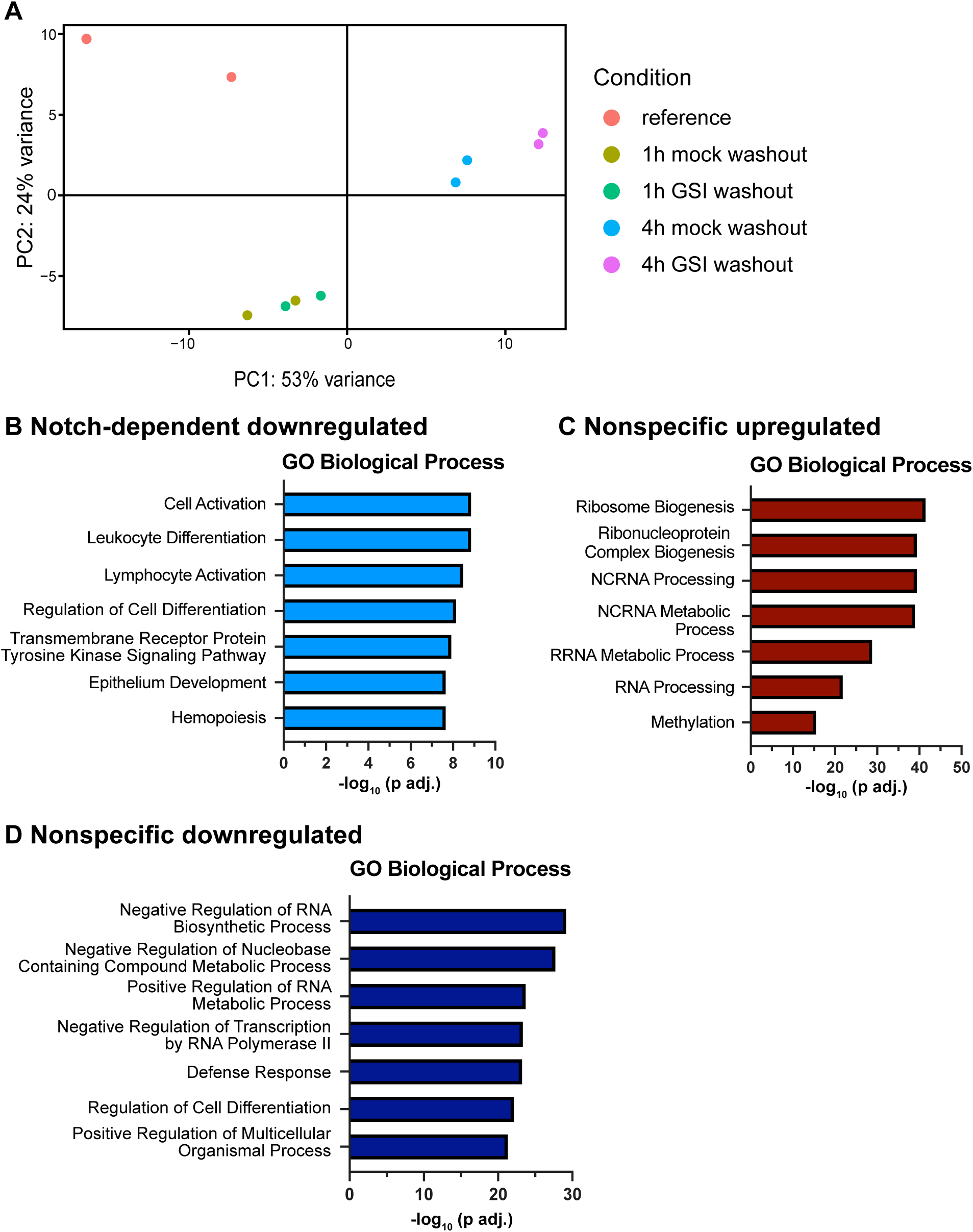
Principal component analysis and gene ontology of the TT-seq experiment, related to Figure 1. A) Principal Component Analysis of TT-seq counts over gene exons for samples shown in Figure1. B-D) Top Gene Ontology – Biological Process enriched terms for genes in the Notch downregulated (B), nonspecific upregulated (C), and nonspecific downregulated (D) groups. P-values are from the hypergeometric test, corrected for

**Figure S2.**
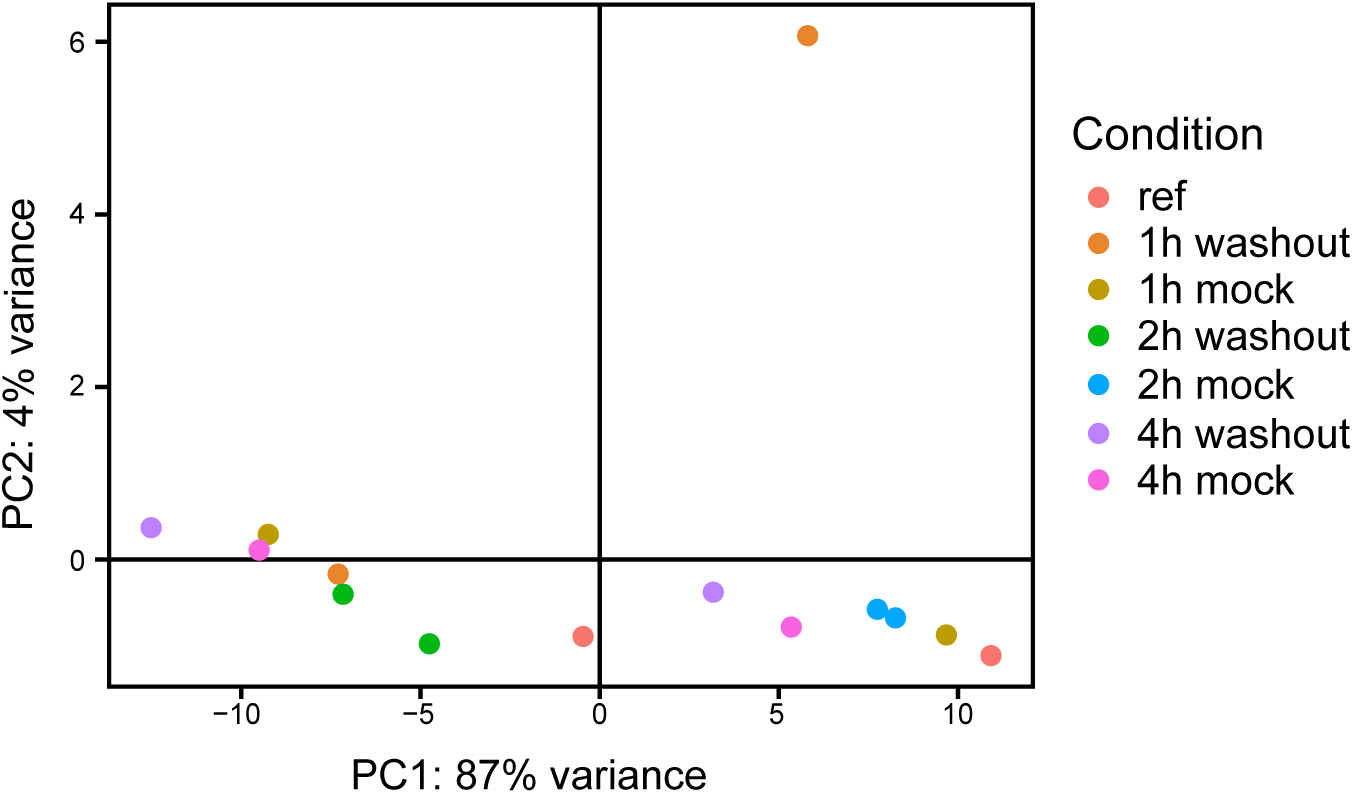
Principal component analysis of ATAC-seq data, related to Figure 2. Principal Component Analysis of ATAC-seq counts over promoter windows for samples in Figure 2.

**Figure S3.**
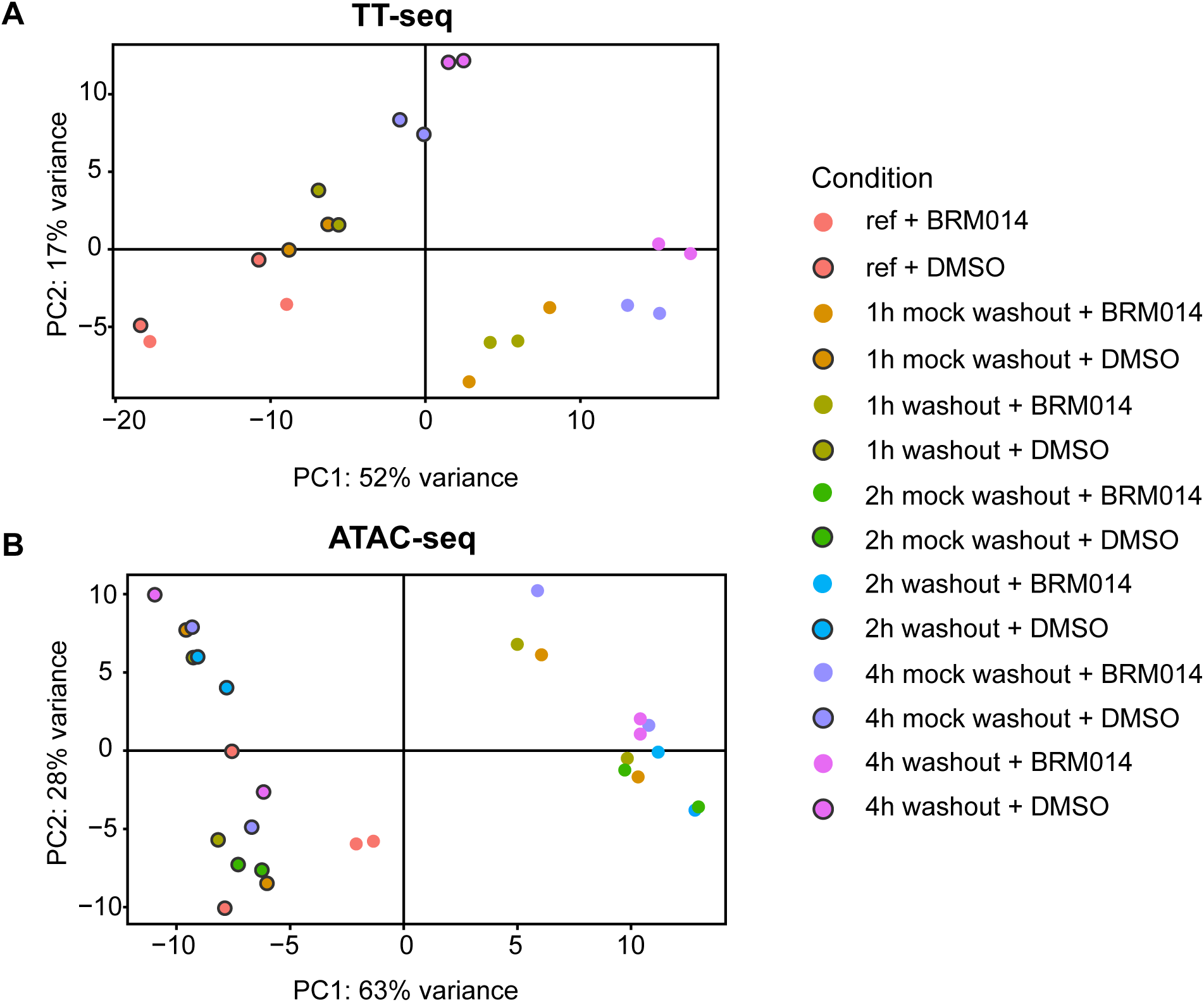
Analysis of chromatin accessibility and gene expression after SWI/SNF inhibition, related to Figure 3. (A) Principal Component Analysis of TT-seq counts over gene exons for samples in Figure 3. (B) Principal Component

**Figure S4.**
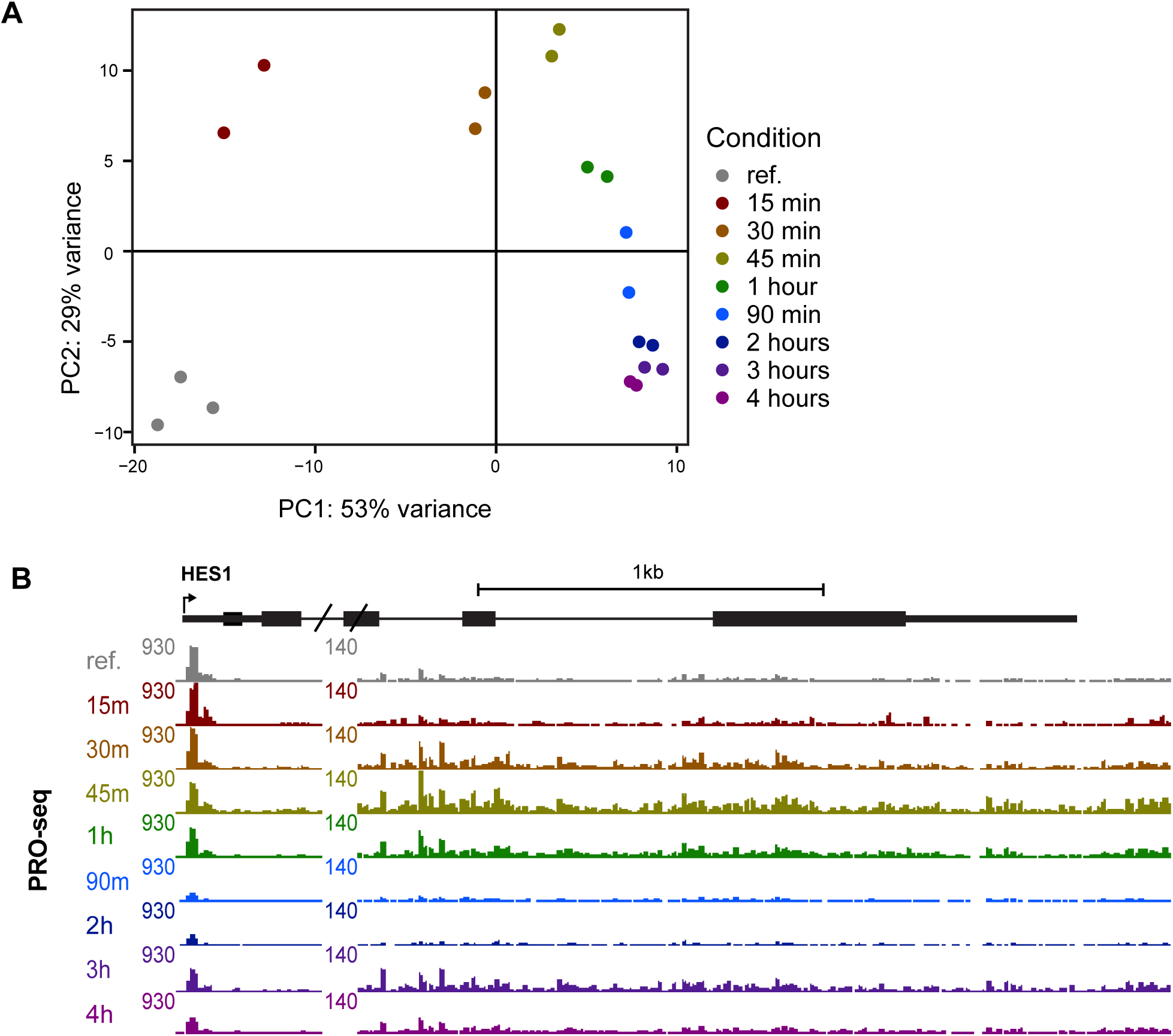
Principal component analysis of PRO-seq data, related to Figure 4. (A) Principal Component Analysis of PRO-seq counts over gene body windows for samples in Figure 4. (B) Genome browser images showing PRO-seq reads for HES1 at all assayed time points. The scale of the browser images is reset at 500 bp downstream of the TSS to allow visualization of the gene body signal.

## REFERENCES

1. Siebel, C., and Lendahl, U. (2017). Notch signaling in development, tissue homeostasis, and disease. Physiol Rev 97, 1235–1294. 10.1152/physrev.00005.2017.

2. Aster, J.C., Pear, W.S., and Blacklow, S.C. (2017). The Varied Roles of Notch in Cancer. Annual Review of Pathology: Mechanisms of Disease 12, 245–275. 10.1146/annurev-pathol-052016-100127.

3. Bray, S.J., and Gomez-Lamarca, M. (2018). Notch after cleavage. Curr Opin Cell Biol 51, 103–109. 10.1016/j.ceb.2017.12.008.

4. Adelman, K., and Lis, J.T. (2012). Promoter-proximal pausing of RNA polymerase II: emerging roles in metazoans. Nat Rev Genet 13, 720–731. 10.1038/nrg3293.

5. Core, L., and Adelman, K. (2019). Promoter-proximal pausing of RNA polymerase II: A nexus of gene regulation. Genes Dev 33, 960–982. 10.1101/gad.325142.119.

6. Pillidge, Z., and Bray, S.J. (2019). SWI / SNF chromatin remodeling controls Notch-responsive enhancer accessibility. EMBO Rep 20, 1–16. 10.15252/embr.201846944.

7. Takeuchi, J.K., Lickert, H., Bisgrove, B.W., Sun, X., Yamamoto, M., Chawengsaksophak, K., Hamada, H., Yost, H.J., Rossant, J., and Bruneau, B.G. (2007). Baf60c is a nuclear Notch signaling component required for the establishment of left – right asymmetry. Proceedings of the National Academy of Sciences 104, 846–851. 10.1073/pnas.0608118104.

8. Yatim, A., Benne, C., Sobhian, B., Laurent-Chabalier, S., Deas, O., Judde, J.G., Lelievre, J.D., Levy, Y., and Benkirane, M. (2012). NOTCH1 Nuclear Interactome Reveals Key Regulators of Its Transcriptional Activity and Oncogenic Function. Mol Cell 48, 445–458. 10.1016/j.molcel.2012.08.022.

9. Martin, A.P., Bradshaw, G.A., Eisert, R.J., Egan, E.D., Tveriakhina, L., Rogers, J.M., Dates, A.N., Scanavachi, G., Aster, J.C., Kirchhausen, T., et al. (2023). A spatiotemporal Notch interaction map from plasma membrane to nucleus. Sci Signal 16.

10. Kadam, S., and Emerson, B.M. (2003). Transcriptional Specificity of Human SWI/SNF BRG1 and BRM Chromatin Remodeling Complexes. Mol Cell 11, 377–389.

11. Gomez-Lamarca, M.J., Falo-Sanjuan, J., Stojnic, R., Abdul Rehman, S., Muresan, L., Jones, M.L., Pillidge, Z., Cerda-Moya, G., Yuan, Z., Baloul, S., et al. (2018). Activation of the Notch Signaling Pathway In Vivo Elicits Changes in CSL Nuclear Dynamics. Dev Cell 44, 611–623.e7. 10.1016/j.devcel.2018.01.020.

12. Skalska, L., Stojnic, R., Li, J., Fischer, B., Cerda-moya, G., Sakai, H., Tajbakhsh, S., Russell, S., Adryan, B., and Bray, S.J. (2015). Chromatin signatures at Notch-regulated enhancers reveal large-scale changes in H3K56ac upon activation. EMBO J 34, 1889–1904. 10.15252/embj.201489923.

13. van den Ameele, J., Cheetham, S.W., Krautz, R., Donovan, A.P.A., Llora-Batlle, O., Yakob, R., and Brand, A.H. (2022). Reduced chromatin accessibility confers resistance to Notch activation. Nat Commun 13. 10.1038/s41467-022-29834-z.

14. Wang, H., Zou, J., Zhao, B., Johannsen, E., Ashworth, T., Wong, H., Pear, W.S., Schug, J., Blacklow, S.C., Arnett, K.L., et al. (2011). Genome-wide analysis reveals conserved and divergent features of Notch1/RBPJ binding in human and murine T-lymphoblastic leukemia cells. Proc Natl Acad Sci U S A 108, 14908–14913. 10.1073/pnas.1109023108.

15. Falo-Sanjuan, J., Lammers, N.C., Garcia, H.G., and Bray, S.J. (2019). Enhancer Priming Enables Fast and Sustained Transcriptional Responses to Notch Signaling. Dev Cell 50, 411–425.e8. 10.1016/j.devcel.2019.07.002.

16. Aster, J.C. (2020). Notch Signaling in Context: Basic and Translational Implications. Trans Am Clin Climatol Assoc 131, 147–156.

17. Pan, L., Lemieux, M.E., Thomas, T., Rogers, J.M., Lipper, C.H., Lee, W., Johnson, C., Sholl, L.M., South, A.P., Marto, J.A., et al. (2020). IER5, a dna damage response gene, is required for notch-mediated induction of squamous cell differentiation. Elife 9, 1–32. 10.7554/ELIFE.58081.

18. Schwalb, B., Michel, M., Zacher, B., Hauf, K.F., Demel, C., Tresch, A., Gagneur, J., and Cramer, P. (2016). TT-seq maps the human transient transcriptome. Science (1979) 352, 1225–1228. 10.1126/science.aad9841.

19. Buenrostro, J.D., Giresi, P.G., Zaba, L.C., Chang, H.Y., and Greenleaf, W.J. (2013). Transposition of native chromatin for fast and sensitive epigenomic profiling of open chromatin, DNA-binding proteins and nucleosome position. Nat Methods 10, 1213–1218. 10.1038/nmeth.2688.

20. Grandi, F.C., Modi, H., Kampman, L., and Corces, M.R. (2022). Chromatin accessibility profiling by ATAC-seq. Nat Protoc 17, 1518–1552. 10.1038/s41596-022-00692-9.

21. Wang, H., Zang, C., Taing, L., Arnett, K.L., Wong, Y.J., Pear, W.S., Blacklow, S.C., Liu, X.S., and Aster, J.C. (2014). NOTCH1-RBPJ complexes drive target gene expression through dynamic interactions with superenhancers. Proceedings of the National Academy of Sciences 111, 705–710. 10.1073/pnas.1315023111.

22. Papillon, J.P.N., Nakajima, K., Adair, C.D., Hempel, J., Jouk, A.O., Karki, R.G., Mathieu, S., Möbitz, H., Ntaganda, R., Smith, T., et al. (2018). Discovery of Orally Active Inhibitors of Brahma Homolog (BRM)/SMARCA2 ATPase Activity for the Treatment of Brahma Related Gene 1 (BRG1)/SMARCA4-Mutant Cancers. J Med Chem 61, 10155–10172. 10.1021/acs.jmedchem.8b01318.

23. Iurlaro, M., Stadler, M.B., Masoni, F., Jagani, Z., Galli, G.G., and Schübeler, D. (2021). Mammalian SWI/SNF continuously restores local accessibility to chromatin. Nat Genet 53, 279–287. 10.1038/s41588-020-00768-w.

24. Schick, S., Grosche, S., Kohl, K.E., Drpic, D., Jaeger, M.G., Marella, N.C., Imrichova, H., Lin, J.M.G., Hofstätter, G., Schuster, M., et al. (2021). Acute BAF perturbation causes immediate changes in chromatin accessibility. Nat Genet 53, 269–278. 10.1038/s41588-021-00777-3.

25. Martin, B.J.E., Ablondi, E.F., Goglia, C., Mimoso, C.A., Espinel-Cabrera, P.R., and Adelman, K. (2023). Global identification of SWI/SNF targets reveals compensation by EP400. Cell 186, 5290–5307.e26. 10.1016/j.cell.2023.10.006.

26. Kwak, H., Fuda, N.J., Core, L.J., and Lis, J.T. (2013). Precise Maps of RNA Polymerase Reveal How Promoters Direct Initiation and Pausing. Science (1979) 339, 950–953.

27. Mahat, D.B., Kwak, H., Booth, G.T., Jonkers, I.H., Danko, C.G., Patel, R.K., Waters, C.T., Munson, K., Core, L.J., and Lis, J.T. (2016). Base-pair-resolution genome-wide mapping of active RNA polymerases using precision nuclear run-on (PRO-seq). Nat Protoc 11, 1455–1476. 10.1038/nprot.2016.086.

28. Hirata, H., Yoshiura, S., Ohtsuka, T., Bessho, Y., Harada, T., Yoshikawa, K., and Kageyama, R. (2002). Oscillatory expression of the bHLH factor Hes1 regulated by a negative feedback loop. Science (1979) 298, 840–843. 10.1126/science.1074560.

29. Gilchrist, D.A., Dos Santos, G., Fargo, D.C., Xie, B., Gao, Y., Li, L., and Adelman, K. (2010). Pausing of RNA polymerase II disrupts DNA-specified nucleosome organization to enable precise gene regulation. Cell 143, 540–551. 10.1016/j.cell.2010.10.004.

30. Heinz, S., Benner, C., Spann, N., Bertolino, E., Lin, Y.C., Laslo, P., Cheng, J.X., Murre, C., Singh, H., and Glass, C.K. (2010). Simple Combinations of Lineage-Determining Transcription Factors Prime cis-Regulatory Elements Required for Macrophage and B Cell Identities. Mol Cell 38, 576–589. 10.1016/j.molcel.2010.05.004.

31. Biddie, S.C., John, S., Sabo, P.J., Thurman, R.E., Johnson, T.A., Schiltz, R.L., Miranda, T.B., Sung, M.H., Trump, S., Lightman, S.L., et al. (2011). Transcription Factor AP1 Potentiates Chromatin Accessibility and Glucocorticoid Receptor Binding. Mol Cell 43, 145–155. 10.1016/j.molcel.2011.06.016.

32. Lefterova, M.I., Steger, D.J., Zhuo, D., Qatanani, M., Mullican, S.E., Tuteja, G., Manduchi, E., Grant, G.R., and Lazar, M.A. (2010). Cell-Specific Determinants of Peroxisome Proliferator-Activated Receptor γ Function in Adipocytes and Macrophages. Mol Cell Biol 30, 2078–2089. 10.1128/mcb.01651-09.

33. Weinmann, A.S., Mitchell, D.M., Sanjabi, S., Bradley, M.N., Hoffmann, A., Liou, H.-C., and Smale, S.T. (2001). Nucleosome remodeling at the IL-12 p40 promoter is a TLR-dependent, Rel-independent event. Nat Immunol 2, 51–57.

34. Saccani, S., Pantano, S., and Natoli, G. (2001). Two Waves of Nuclear Factor κB Recruitment to Target Promoters. J. Exp. Med 193, 1351–1359.

35. Utley, R.T., Côté, J., Owen-Hughes, T., and Workman, J.L. (1997). SWI/SNF stimulates the formation of disparate activator-nucleosome complexes but is partially redundant with cooperative binding. Journal of Biological Chemistry 272, 12642–12649. 10.1074/jbc.272.19.12642.

36. Hargreaves, D.C., Horng, T., and Medzhitov, R. (2009). Control of Inducible Gene Expression by Signal-Dependent Transcriptional Elongation. Cell 138, 129–145. 10.1016/j.cell.2009.05.047.

37. Adelman, K., Kennedy, M.A., Nechaev, S., Gilchrist, D.A., Muse, G.W., Chinenov, Y., and Rogatsky, I. (2009). Immediate mediators of the inflammatory response are poised for gene activation through RNA polymerase II stalling. Proceedings of the National Academy of Sciences 106, 18207–18212.

38. Weng, A.P., Millholland, J.M., Yashiro-Ohtani, Y., Arcangeli, M.L., Lau, A., Wai, C., Del Bianco, C., Rodriguez, C.G., Sai, H., Tobias, J., et al. (2006). C-Myc is an important direct target of Notch1 in T-cell acute lymphoblastic leukemia/lymphoma. Genes Dev 20, 2096–2109. 10.1101/gad.1450406.8.

39. Lagha, M., Bothma, J.P., Esposito, E., Ng, S., Stefanik, L., Tsui, C., Johnston, J., Chen, K., Gilmour, D.S., Zeitlinger, J., et al. (2013). Paused Pol II coordinates tissue morphogenesis in the drosophila embryo. Cell 153, 976. 10.1016/j.cell.2013.04.045.

40. Boettiger, A.N., and Levine, M. (2009). Synchronous and Stochastic Patterns of Gene Activation in the Drosophila Embryo. Science (1979) 325, 471–473. 10.1126/science.1098641.

41. Fryer, C.J., Lamar, E., Turbachova, I., Kintner, C., and Jones, K.A. (2002). Mastermind mediates chromatin-specific transcription and turnover of the notch enhancer complex. Genes Dev 16, 1397–1411. 10.1101/gad.991602.

42. Rogers, J.M., Guo, B., Egan, E.D., Aster, J.C., Adelman, K., and Blacklow, S.C. (2020). MAML1-Dependent Notch-Responsive Genes Exhibit Differing Cofactor Requirements for Transcriptional Activation. Mol Cell Biol 40, 1–11. 10.1128/mcb.00014-20.

43. Liu, K., Shen, D., Shen, J., Gao, S.M., Li, B., Wong, C., Feng, W., and Song, Y. (2017). The Super Elongation Complex Drives Neural Stem Cell Fate Commitment. Dev Cell 40, 537–551.e6. 10.1016/j.devcel.2017.02.022.

44. Grandi, F.C., Modi, H., Kampman, L., and Corces, M.R. (2022). Chromatin accessibility profiling by ATAC-seq. Nat Protoc 17, 1518–1552. 10.1038/s41596-022-00692-9.

45. Langmead, B., Trapnell, C., Pop, M., and Salzberg, S.L. (2009). Ultrafast and memory-efficient alignment of short DNA sequences to the human genome. Genome Biol 10. 10.1186/gb-2009-10-3-r25.

46. Dobin, A., Davis, C.A., Schlesinger, F., Drenkow, J., Zaleski, C., Jha, S., Batut, P., Chaisson, M., and Gingeras, T.R. (2013). STAR: Ultrafast universal RNA-seq aligner. Bioinformatics 29, 15–21. 10.1093/bioinformatics/bts635.

47. Love, M.I., Huber, W., and Anders, S. (2014). Moderated estimation of fold change and dispersion for RNA-seq data with DESeq2. Genome Biol 15. 10.1186/s13059-014-0550-8.

48. Langmead, B., and Salzberg, S.L. (2012). Fast gapped-read alignment with Bowtie 2. Nat Methods 9, 357–359. 10.1038/nmeth.1923.

49. Kent, W.J., Zweig, A.S., Barber, G., Hinrichs, A.S., and Karolchik, D. (2010). BigWig and BigBed: Enabling browsing of large distributed datasets. Bioinformatics 26, 2204–2207. 10.1093/bioinformatics/btq351.

50. Liao, Y., Smyth, G.K., and Shi, W. (2014). FeatureCounts: An efficient general purpose program for assigning sequence reads to genomic features. Bioinformatics 30, 923–930. 10.1093/bioinformatics/btt656.

51. Danko, C.G., Hyland, S.L., Core, L.J., Martins, A.L., Waters, C.T., Lee, H.W., Cheung, V.G., Kraus, W.L., Lis, J.T., and Siepel, A. (2015). Identification of active transcriptional regulatory elements from GRO-seq data. Nat Methods 12, 433–438. 10.1038/nmeth.3329.

52. Liberzon, A., Subramanian, A., Pinchback, R., Thorvaldsdóttir, H., Tamayo, P., and Mesirov, J.P. (2011). Molecular signatures database (MsigDB) 3.0. Bioinformatics 27, 1739–1740. 10.1093/bioinformatics/btr260.

## Supplemental Methods References

53. Duffy, E.E., Rutenberg-Schoenberg, M., Stark, C.D., Kitchen, R.R., Gerstein, M.B., and Simon, M.D. (2015). Tracking Distinct RNA Populations Using Efficient and Reversible Covalent Chemistry. Mol Cell 59, 858–866. 10.1016/j.molcel.2015.07.023.

54. Dölken, L., Ruzsics, Z., Rädle, B., Friedel, C.C., Zimmer, R., Mages, J., Hoffmann, R., Dickinson, P., Forster, T., Ghazal, P., et al. (2008). High-resolution gene expression profiling for simultaneous kinetic parameter analysis of RNA synthesis and decay. RNA 14, 1959–1972. 10.1261/rna.1136108.

55. Reimer, K.A., Mimoso, C.A., Adelman, K., and Neugebauer, K.M. (2021). Co-transcriptional splicing regulates 3′ end cleavage during mammalian erythropoiesis. Mol Cell 81, 998–1012.e7. 10.1016/j.molcel.2020.12.018.

56. Martin, M. (2011). Cutadapt removes adapter sequences from high-throughput sequencing reads. EMBnet J 17, 10–12.

57. Li, H., Handsaker, B., Wysoker, A., Fennell, T., Ruan, J., Homer, N., Marth, G., Abecasis, G., and Durbin, R. (2009). The Sequence Alignment/Map format and SAMtools. Bioinformatics 25, 2078–2079. 10.1093/bioinformatics/btp352.

